# The Current Landscape of Plasma Proteomics: Technical Advances, Biological Insights, and Biomarker Discovery

**DOI:** 10.1101/2025.02.14.638375

**Authors:** Douglas Y. Kirsher, Shreya Chand, Aron Phong, Bich Nguyen, Balazs G. Szoke, Sara Ahadi

**Affiliations:** Alkahest Inc. San Carlos, CA

**Author notes:** Corresponding author Correspondence to: Sara Ahadi.

## Abstract

Plasma is a rich source of biomolecules, including proteins, that reflect both health and disease. Due to their key roles in biological processes, proteins hold significant potential as biomarkers, fueling the rise of plasma proteome profiling in recent years. Despite widespread adoption, few studies have directly compared different plasma proteomics platforms, particularly those using mass spectrometry. Our study provides a comprehensive comparison of seven platforms across three leading technologies - SomaLogic, Olink, and Mass Spectrometry (MS) - including affinity-based approaches and various MS techniques, covering over 13,000 proteins. By applying these methods to the same cohort, we assess their performance, revealing key differences and complementary strengths. Our findings offer valuable insights for researchers, highlighting trade-offs in coverage and their implications for biomarker discovery and clinical applications. This study serves as an essential resource, offering both technical evaluation and biological insights to support the development of novel diagnostics and therapeutics through plasma proteomics.

## Introduction

Plasma is a key component of blood, containing a diverse array of molecules, including proteins, nucleic acids (DNA and RNA), lipids, small molecule metabolites, and electrolytes. Many of these molecules are essential for maintaining various physiological processes within the body.^1^ Changes in their concentrations can signal or drive underlying health issues, making plasma a valuable resource for discovering biomarkers or therapeutic targets for diseases. One of the main benefits of using plasma for this purpose is the relative ease of collection. Plasma samples are inexpensive to collect and minimally invasive to obtain, and the procedure is well-tolerated by patients, making it ideal for routine use in clinical settings. This simplicity enables longitudinal collection, which provides richer data for biomarker discovery, target identification, and insights into system biology and disease progression.

While plasma is a promising source of various biomarkers, proteins are particularly significant due to their essential roles in biological processes and disease mechanisms.^2,3^ Given their direct involvement in cellular functions and potential for therapeutic targeting, proteins often emerge as key indicators of health and disease states.^4^ Moreover, analysing plasma protein levels alongside genetic and phenotypic information can create a comprehensive picture that enhances our understanding of health status and improves disease management.

However, utilizing the plasma proteome as a source of biomarkers presents several challenges.^3,5,6,7^ The plasma proteome spans a wide dynamic range, with some biomarkers present at very low concentrations, complicating their detection and robust quantification. Additionally, data interpretation requires careful consideration of confounding factors such as age,^8,9^ sex,^9^ BMI,^9,10^ and fasting status, besides technical aspects^6,11^ such as storage duration^12,13,14^ and temperature,^13,14^ and blood collection factors such as time of the day, the use of different anticoagulants,^15^ and blood processing protocol.^13,14^ Studies have also indicated that while individual proteome profiles tend to remain stable over time, there is significant variability between individuals.^7^ These challenges emphasize the need for robust and complementary methodologies, along with a thorough understanding of the data, to effectively leverage the plasma proteome for biomarker and therapeutics development.

Currently, two primary approaches are used to analyze plasma proteins: affinity-based techniques and mass spectrometry (MS) based methods. Affinity-based platforms, such as SomaLogic’s and Olink’s, use binding probes, aptamers or antibodies, respectively, to detect proteins. In contrast, MS methods typically measure proteolytic peptides of proteins through a bottom-up approach. To address the dynamic range challenge, MS workflows may include high-abundance protein depletion,^16^ protein pre-fractionation,^17^ protein precipitation,^18^ ion mobility-based separation,^16^ or protein enrichment via beads^19^ or nanoparticles.^20^

Each method measures distinct characteristics of plasma proteins and has its own unique advantages and disadvantages. SomaLogic and Olink utilize targeted assays that rely on the pre-selection of targets. These techniques allow for high-throughput measurements and multiplexing, enabling the analysis of thousands of proteins from small sample volumes. However, affinity binding techniques can be non-specific, and their accuracy is often dependent on the epitope and the matrix. SomaLogic’s assays rely on the unique engagement of a specific binder for each target protein, which can introduce bias based on the matrix and environment in which the protein is measured. Publicly available information on the binding reagents used in SomaScan assays (SOMAmers) and their availability for pull-down assays can help improve our understanding of the specificity of each aptamer.^21^ Olink’s proximity extension assays mitigate this specificity issue by requiring two different antibodies to bind to the target protein in close proximity.^22^ In contrast, MS-based proteomics is unbiased and can be performed via untargeted or targeted approaches. MS-based assays typically measure multiple peptides of one protein and are capable of identification of post translational modifications (PTMs) and isoforms of proteins, offering unique specificity in protein identification.^22,23^ However, its limited depth makes it challenging to use for plasma, which has a dynamic range spanning 10 orders of magnitude. Despite this limitation, advancements in mass spectrometry have improved proteome coverage and throughput, allowing for the identification and quantification of a broader range of proteins, including clinically relevant ones present in lower abundance.^24^

This study offers a comprehensive comparison of state-of-the-art plasma proteome analysis techniques, including SomaLogic, Olink, and MS-based methods, by applying each technology to the same cohort of plasma samples. This allowed for a detailed analysis of each platform, identifying their advantages and disadvantages. We examined multiple versions of affinity-based platforms (SomaScan 11K, SomaScan 7K, Olink Explore 3072 and Explore HT) alongside discovery MS workflows, including the nanoparticle-based Seer Proteograph™ platform, and a high abundance protein depletion-based workflow - Biognosys TrueDiscovery™ platform, both utilizing Data-Independent Acquisition (DIA) to generate deep, unbiased MS data. Additionally, we incorporated a targeted MS workflow, SureQuant™ Internal Standard Triggered - Parallel Reaction Monitoring, as a "gold standard" MS platform of high reliability and absolute quantification due to the use of internal standards and optimized detection. To the best of our knowledge, this represents the largest number of plasma proteomic technologies compared using a single cohort. By juxtaposing these methods, we aim to provide a detailed technical evaluation and biological insight that will serve as a valuable resource for researchers in plasma proteomics. Understanding the unique characteristics of each platform will assist investigators in selecting the most appropriate method for their research objectives and guide future biomarker discovery efforts.

## Results

### Cohort and Platforms

In our analysis, we obtained plasma protein profiles from a cohort of 78 individuals, maintaining an equal sex ratio of 1:1 (male to female) consisting of 40 aged (55-65 years old) and 38 young (18-22 years old) individuals. Summary of cohort demographics is listed in Supplementary Table S1. Plasma samples were collected via plasmapheresis and analysed using seven proteomic platforms, as shown in Figure 1. For clarity, we will refer to Olink Explore 3072 and Olink Explore HT as Olink 3K and Olink 5K, respectively, for the remainder of this discussion. Similarly, MS-based workflows will be designated as MS-Nanoparticle (Seer Proteograph™ XT), MS-HAP Depletion (Biognosys TrueDiscovery™ platform using high-abundance protein depletion), and MS-IS Targeted (SureQuant™ Internal Standard Triggered - Parallel Reaction Monitoring). SomaLogic 11K includes 11,083 assays, with 10,776 human protein assays targeting 9,852 unique proteins, corresponding to 9,645 distinct UniProt IDs. SomaLogic 7K includes 7,596 assays, with 7,288 human protein assays targeting 6,467 unique proteins, corresponding to 6,401 distinct UniProt IDs. Olink 5K and 3K assays target 5,416 and 2,925 unique human proteins, respectively. In our study, the number of proteins identified by MS-based platforms were 5,943 in MS-Nanoparticle, 3,575 in MS-HAP Depletion and 551 MS-IS Targeted. Each quantified 68,527, 42,581, and 766 peptides, respectively. A complete list of the proteins identified in this study, including UniProt IDs, can be found in Supplementary Table S2.

**Figure 1:**
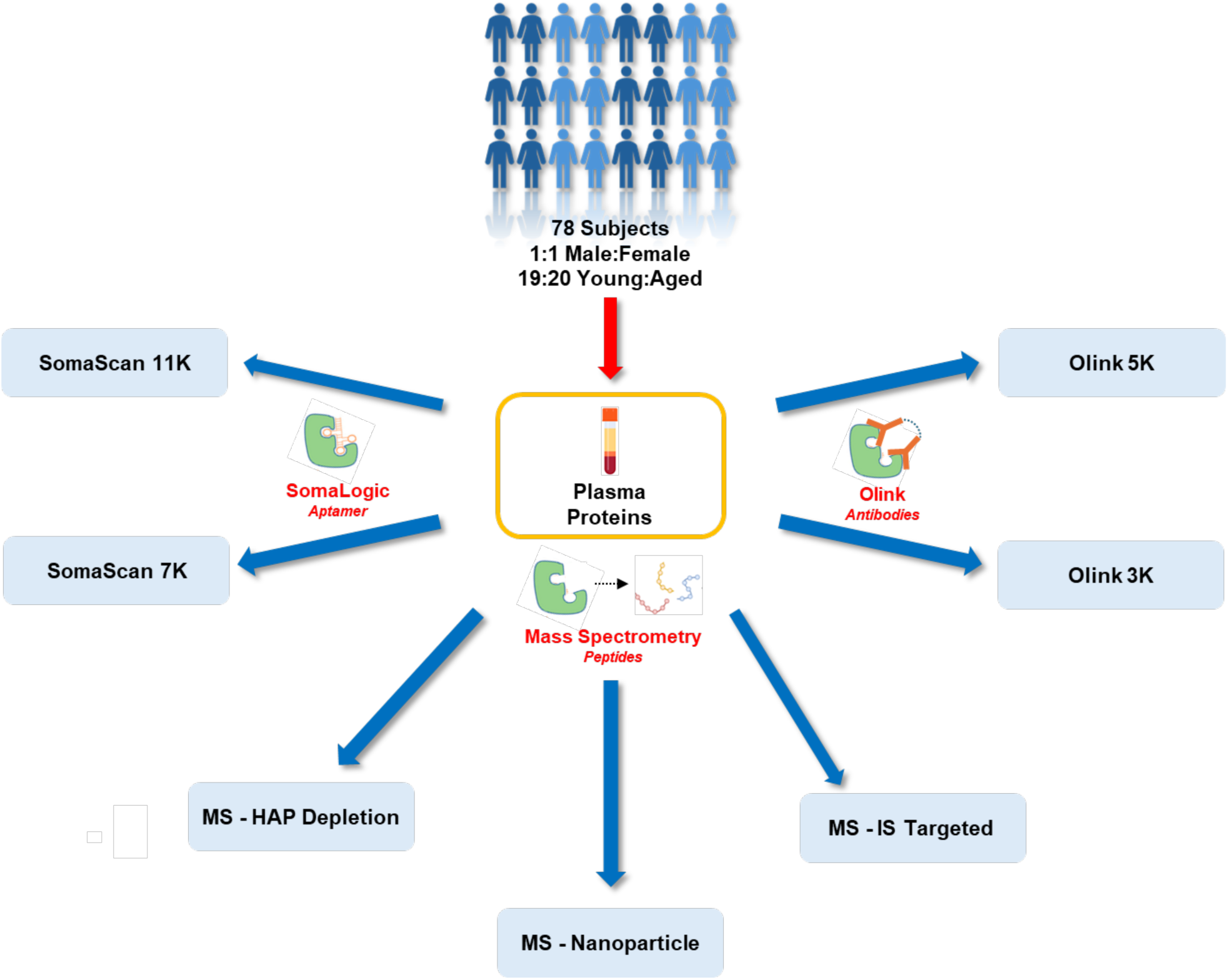
Study Overview. Plasma samples collected via plasmapheresis from aged (n=40) and young (n=38) subjects were analyzed using seven proteomic platforms: SomaScan 11K, SomaScan 7K, Olink 5K (Olink Explore HT), Olink 3K (Olink Explore 3072), MS-HAP Depletion (high-abundance protein depletion), MS-Nanoparticle (Seer Proteograph™ XT) and MS-IS Targeted (SureQuant™ Internal Standard Triggered - Parallel Reaction Monitoring).

### Technical Assessments

Across all seven platforms, we identified a total of 13,007 unique plasma proteins, each represented by a unique Uniprot ID, in our healthy plasma samples. As illustrated in Figure 2A, the SomaScan 11K and SomaScan 7K platforms provided the most comprehensive proteomic coverage, detecting 9,645 and 6,401 proteins, respectively. MS-Nanoparticle followed with 5,943 unique proteins. Each platform contributed a set of exclusive proteins that were not identified by the others. The SomaLogic platforms contributed the largest number of exclusive proteins, with SomaScan 11K identifying 3,610 unique proteins and SomaScan 7K identifying 1,954 unique proteins compared to Olink and MS platforms. The MS-Nanoparticle platform contributed 1,036 exclusive proteins.

**Figure 2:**
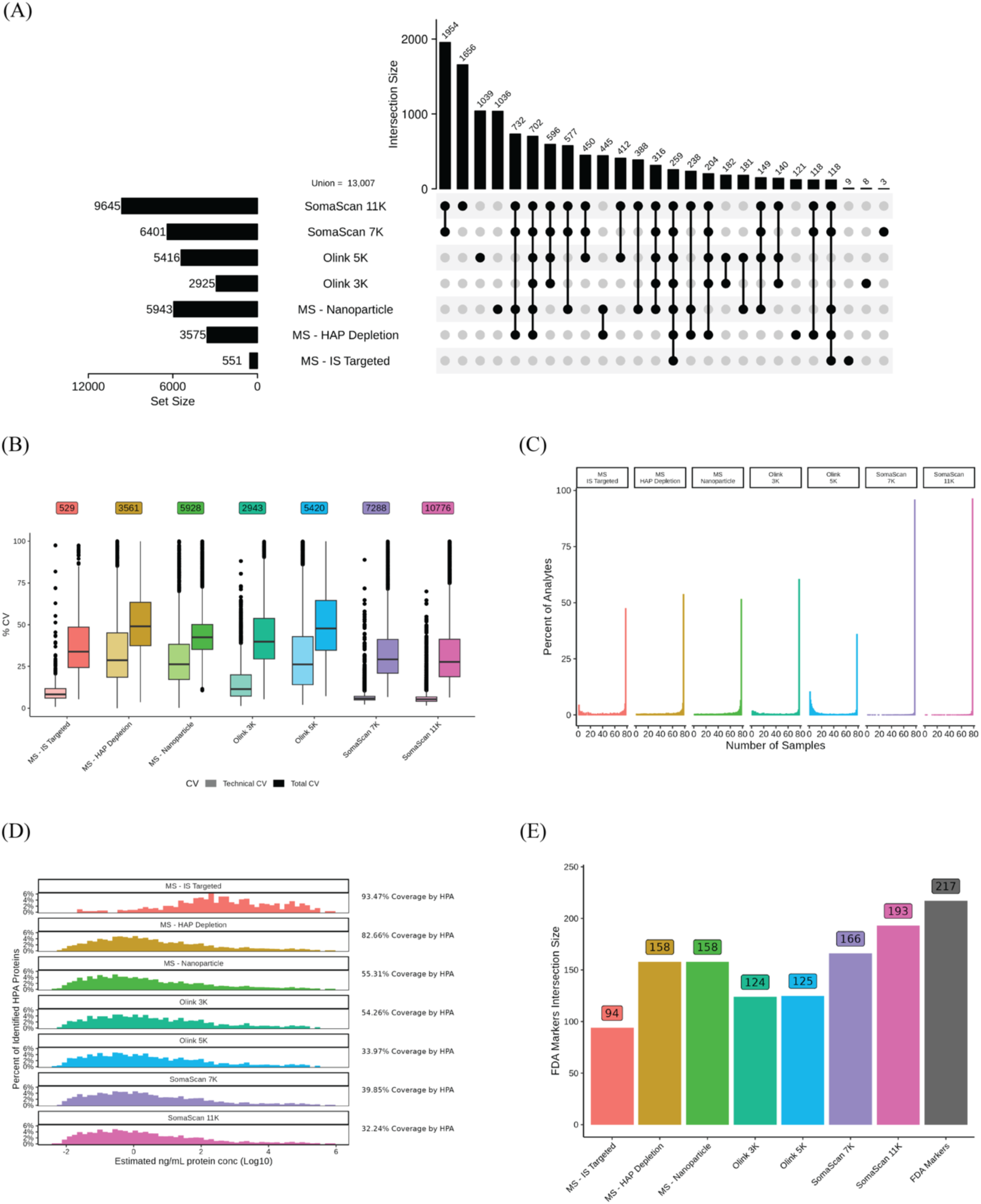
QC metrics for platform measurements. (A) UpSet plot showing intersection of identified proteins from each platform. Only intersection sizes of greater than 100 are shown. 254 shared proteins are quantifiable across all platforms. (B) Technical and total CVs shown for each analyte on each platform. Technical CVs were calculated using technical replicates of n = 2 or n = 3, depending on the platform. Total CVs were calculated using the shared 78 subjects. (C) Percentage of analytes that were detected in each number of samples. MS data was filtered based on 2/3rds data completeness from each age group. Affinity based platform data was filtered based on appropriate LOD. (D) The detected proteins present in the HPA dataset were binned according to the estimated concentrations from the HPA dataset. The percentage make-up of proteins belonging to these bins are plotted. (E) The number of FDA approved protein biomarkers identified by each platform.

Among the seven platforms, there was an overlap of 259 proteins, for which we have absolute quantitation values. This overlap includes MS-IS Targeted data, which was used as a reference to evaluate the other platforms. Excluding MS-IS Targeted, the remaining six platforms detected 702 shared proteins.

To assess the variability in quantification across platforms, technical replicates were utilized. As shown in Figure 2B, SomaScan exhibited the highest precision among all platforms, with the lowest technical coefficients of variation (CV). The median CVs for SomaScan 11K and 7K were 5.3% and 5.8%, respectively, indicating that the addition of over 3,000 assays to the latest version of SomaScan has not compromised the platform’s excellent precision. In contrast, the median CV for Olink 5K was more than twice of that of Olink 3K, at 26.8% and 11.4%, respectively. There have been many technical assessments of SomaScan and Olink assays in literature in the past few years ^24,25,26,27,28^. Our CV results are consistent with these reports and observations. For the MS-based platforms, the technical CVs of discovery-based approaches were higher than those for SomaScan 11K, SomaScan 7K and Olink 3K, with median CVs of 26.4% for MS-Nanoparticle and 29.8% for MS-HAP Depletion. In contrast to discovery MS platforms, MS-IS Targeted had a median CV of 8.3%, due to the optimized targeted analysis of the proteins and the use of internal controls for each analyte. This same trend was also observed when we restricted the results to analytes that were observed across all platforms (Supplementary Figure 1). The median CVs of Olink 5K and 3K were again higher than those of SomaScan 11K and 7K. The MS-based platforms continued to show higher CVs than affinity platforms, except for MS-IS Targeted that had comparable precision to the affinity platforms (Supplementary Table S2). Filtering to include two-thirds of measurements that were above the platform-specific estimated Limit of Detection (eLOD) for affinity-based assays, or to those detected in at least two-thirds of the samples for MS-based platforms, resulted in subtle CV changes for some platforms but more pronounced changes for others (Supplementary Figure 1). The most notable change occurred for Olink 5K, where limiting data to the measurements that were above eLOD for healthy plasma, improved the CV from 26.8% to 12.4%. It is important to note, however, that this was also accompanied by a 40% reduction in the number of analytes used to calculate the CVs after filtering (Supplementary Table S3).

As expected, the median total CV values across all samples, incorporating both biological and technical variability, were higher than the corresponding technical CVs for all platforms. The gap between technical and biological CV varied widely among the different proteomic approaches. For the two SomaScan platforms, Olink 3K and MS-IS Targeted, technical CV has little to no interference with biological variability, while for the two MS discovery approaches and Olink 5K, the technical variability accounted for a much larger portion of the total observed variability.

For the affinity probe-based platforms, SomaScan and Olink, we carried out a simple linearity assessment by diluting pooled plasma samples 3 and 9 times and checking for the linearity of the measured protein signals. Pearson correlation coefficient (r) was calculated for the dilution data and used to characterize the linearity of each protein assay (Supplementary Figure 2). Our results showed that 97% of all SomaScan assays (both 7K and 11K) detected normal plasma protein levels in their linear range (r > 0.9), while the same measure was found to be 42% for the assays of the Olink 5K platform. Limiting the analysis to assays in which all 3 dilutions yielded values above eLOD, we found that high proportion of the Olink assays also showed linear behaviour (85% of assays with r > 0.9).

To assess data completeness across different platforms, we examined the number of missing values for each and plotted the distribution of data completeness in Figure 2C. For MS data, missing values were defined as either a recorded value of 0 or failure to be detected and for SomaLogic and Olink platforms, missing values were those falling below the platform-specific estimated Limit of Detection (eLOD). Our analysis revealed that SomaLogic had the highest data completeness, with SomaScan 11K and 7K showing 96.2% and 95.8% completeness, respectively, across all 78 samples. The Olink 3K platform followed with 60.3% completeness, and MS-HAP Depletion had 53.6% completeness. Notably, the latest version of Olink platform, 5K, had significantly lower data completeness at 35.9%, compared to the previous 3K version at 60.3%. This finding is consistent with the recent literature on comparison between affinity platforms.^28^

We visualized the abundance of proteins identified in each platform by plotting their distribution based on estimated concentrations of approximately 4,392 plasma proteins from the Human Proteome Atlas (HPA, https://v21.proteinatlas.org/humanproteome/blood+protein). As shown in Figure 2D, all platforms, except MS-Targeted, identified analytes across a wide concentration range (10^5^ - 10^-2^ ng/mL), with a strikingly similar concentration distribution for the proteins overlapping between each platform and the HPA. This suggests that technological advancements in mass spectrometry, such as depletion or enrichment methods, have significantly expanded the MS coverage of the plasma proteome, to levels similar to that by affinity methods. However, it is important to note that the HPA dataset is limited, and this comparison does not consider a large number of proteins detected by only affinity-based platforms which are not included in the HPA dataset.

To assess the clinical utility of each platform, we examined their coverage of known protein biomarkers in human plasma approved by the U.S. Food and Drug Administration (FDA).^29^ This is illustrated in Figure 2E, which shows the distribution of circulating biomarkers across platforms using a previously published list.^30^ SomaLogic platforms demonstrated the highest coverage of these biomarkers. Specifically, 11K and 7K covered 88% and 76% of FDA-approved biomarkers, respectively. This was followed by the discovery MS methods (73%) and Olink (57%). Although MS-IS Targeted had the lowest coverage at 43%, this still represented substantial detection of the 217 FDA-approved markers, considering the relatively small number of total proteins identified using this platform.

### Correlations Between Shared Proteins Among Platforms

With the increase in depth of proteomics platforms, the number of identified proteins covered by the platforms and overlap proteins among all increase and correlation between overlapped proteins can be indicative of their similarities and contrasts in identification and quantitation of proteins. Figure 3A illustrates the number of overlapping proteins between different platforms. The largest overlap between inter-proteomic technologies is 3978 proteins between SomaScan 11K and MS-Nanoparticle, followed by 3720 proteins between Somascan11K and Olink 5K.

**Figure 3:**
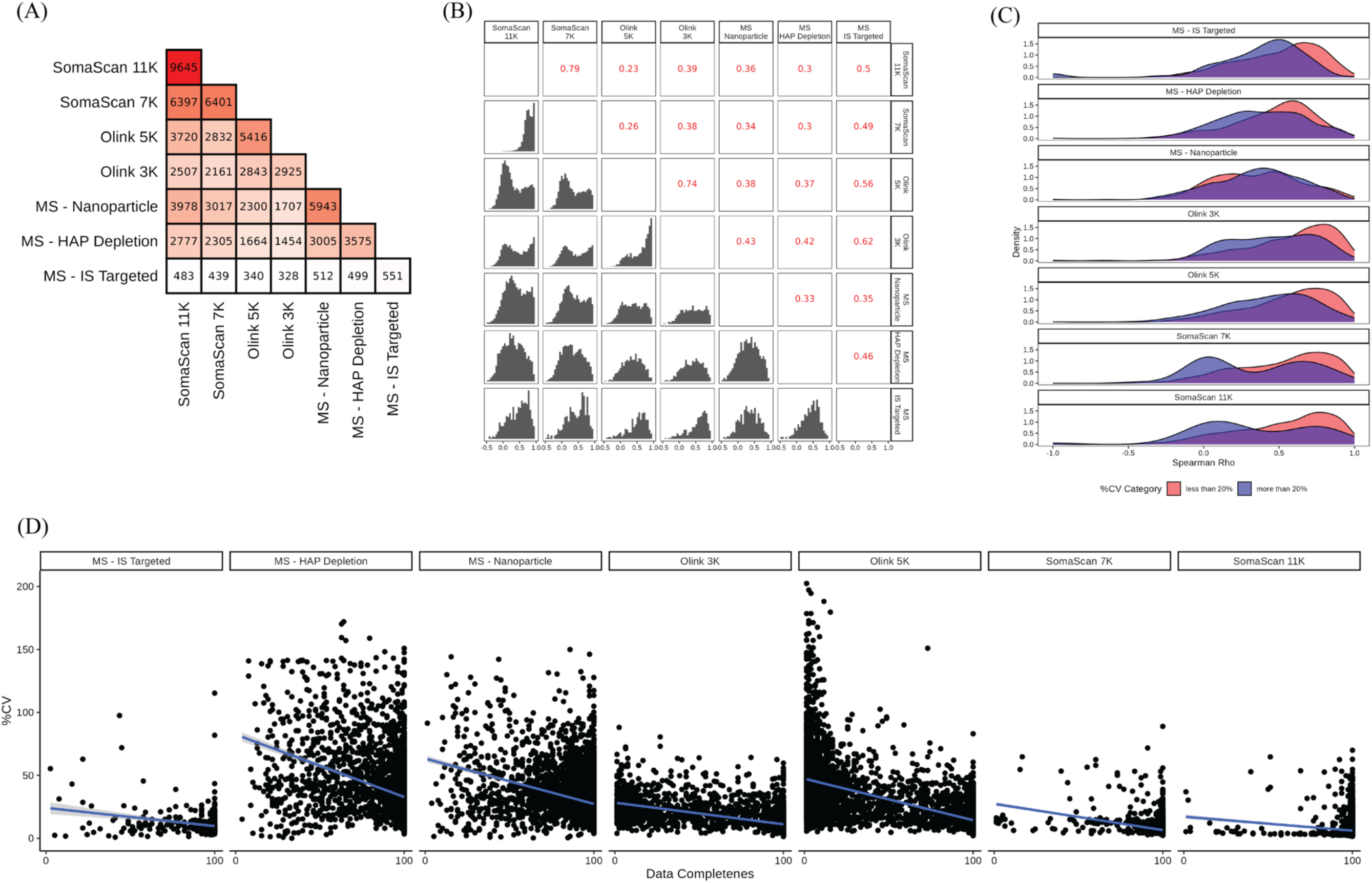
Inter-platform comparisons. (A) Intersection sizes of detected proteins between each pair of platforms. (B) Histogram showing the distribution of Spearman Rho correlation coefficient calculated on a per-protein basis. The median Spearman Rho correlation coefficient is shown in red. (C) Density plot of the Spearman Rho correlation coefficient between a single platform and all other platforms, for the shared 254 proteins across all platforms. The density plot is colored by technical CV of the baseline platform. (D) Percent CV versus Data Completeness for each analyte across each of the platforms.

To further investigate these intersections, we analyzed the correlation of protein intensities across the various platforms. Spearman correlation values for unique protein assays shared between platforms are plotted in Figure 3B and Supplementary Table S4. As expected, two versions of each affinity platform correlated highly with each other, SomaScan 11K and 7K (0.79), followed by Olink 5K and 3K (0.74). When comparing Olink 5K and 3K with other platforms, 5K showed poorer correlation than 3K, while SomaScan 11K and 7K had roughly the same median correlation with other platforms. Among the MS technologies, MS-IS Targeted demonstrated strongest correlations with other platforms (Spearman correlation of 0.35, 0.46, 0.49, 0.5, 0.56 and 0.62 with MS-Nanoparticle, MS-HAP Depletion, SomaLogic 7K, SomaLogic 11K, Olink 5K and Olink 3K respectively). Because of high specificity of targeted MS with internal standard approach, we have observed higher correlations between MS-IS Targeted and other platforms. Notably, MS-IS Targeted showed a correlation of 0.62 with Olink 3K, which was the highest correlation between any of the affinity based and MS based technologies compared in this study. High correlation with MS-IS Targeted, as a reference for assessing other platforms, indicates higher specificity of the measurements in Olink 3K, which has also been reported in literature via *cis*-pQTL validations of affinity assays.^31^ Additionally, all correlations except MS-Nanoparticle exhibited a bimodal pattern, which was more pronounced in some overlaps than in others. This suggests the presence of two groups of proteins with distinct distributions of high and low correlations. Pairwise correlations between each two platforms are presented in Supplementary Figure 3 and 4.

To better understand the reasons behind bimodal distribution of platforms’ pairwise correlations, we focused on the 259 proteins shared by all platforms. The lack of correlation between platforms can be related to several factors, most likely differences in identification specificity and poor quantification precision. In Figure 3C, we examined the relationship between the technical CVs of these proteins within each platform and their correlation with all other platforms. Proteins were categorized into two groups based on their CV values: less than 20%, and higher than 20%. Figure 3C displays distribution of Spearman correlation values for each CV category. This plot revealed that proteins with CVs less than 20% tend to have higher correlations with other platforms. Conversely, proteins with CVs higher than 20% showed a shift towards lower correlations with other platforms. This trend is true for all platforms, except for MS-Nanoparticle, where there is no clear distinction in correlation values with technical CVs of proteins.

Supplementary Figure 5 presents three example proteins with different correlation patterns across platforms. P11226 (MBL2), for example, is one of the proteins that is highly correlated among all platforms, with Spearman correlation values ranging from 0.988-0.863. For some proteins, correlations are high between affinity platforms, but not with the discovery MS platforms. P07359 (GP1BA) illustrates this trend, showing high correlation values (0.931-0.785) among affinity platforms and low correlation values among MS platforms (0.244-0.098). In this case, MS-IS Targeted is well correlated with affinity platforms, indicating the accuracy of their measurements (0.81-0.759). P01008 (SERPINC1), on the other hand, is an example of a protein whose measurements do not correlate between any of the platforms, even showing poor correlation between two different versions of SomaScan and Olink platforms (0.495-0.186).

Lack of correlation may be due to proteoform selectivity differences between the corresponding assays of the different platforms. SomaScan 11K platform include two or more aptamers for 981 unique Uniprot IDs, of which 175 have multiple SOMAmer target IDs. Correlation between assays using aptamers representing one SOMAmer target ID has median Spearman Rho correlation value of 0.54. The distribution of Rho is bimodal, however, with one group of assays showing excellent correlation, and a second group showing weak correlation (Rho ∼ 0.4) between the assays targeting the same protein by different SOMAmers. (Supplementary Figure 6). Identifying exact target of each aptamer requires experimental approaches such as pull-down assays and orthogonal characterization of the pulled-down protein(s).

We also examined the connection between CV and data completeness for each platform. Figure 3D is a scatterplot showing the percentage of data completeness versus CV, illustrating a clear negative correlation between the two. This correlation was more pronounced in MS-HAP Depletion, MS-Nanoparticle and Olink platforms, which also had the highest number of missing values. A similar correlation between CV and data completeness has been reported for Olink 5K, where CV was strongly inversely correlated (r = - 0.77) with protein detectability, consistent with our findings.^28^

### Biological Relevance and Variance Analysis of Identified Proteins

To capture the biology covered by these proteomics platforms, we looked into protein classes defined by the PANTHER Classification System. Identified proteins from each platform were categorized into protein classes. Figure 4A shows a heatmap representing the distribution of PANTHER protein classes across different platforms. There is a vast difference in protein class coverage of these platforms, primarily due to the number of covered proteins or targeted/untargeted nature of each platform (Supplementary Table S5). As expected, SomaScan 11K covered the highest number of protein classes compared to the other platforms. Common protein classes across all platforms, with at least 30% representation include apolipoprotein, complement component, protease inhibitor, and surfactant protein classes. The Runt transcription factor class was uniquely characterized by SomaScan platforms, whereas mitochondrial carrier proteins was uniquely characterized by SomaScan 11K and MS-Nanoparticle platforms. While DNA methyltransferase was uniquely characterized by SomaScan and MS-Nanoparticle platforms, DNA photolyase was uniquely characterized by SomaScan 11K with 100% representation (class size of 2 proteins). Further, SomaScan 11K was able to characterize seven additional protein classes (DNA ligase, DNA photolyase, adenylate cyclase, centromere DNA-binding protein, deacetylase, mitochondrial carrier protein, tubulin), compared to SomaScan 7K. Olink 5K characterized 13 additional protein classes (DNA ligase, MADS box transcription factor, RNA methyltransferase, adenylate cyclase, amino acid transporter, centromere DNA-binding protein, gene-specific transcriptional regulator, glucosidase, mRNA capping factor, mRNA polyadenylation factor, primase, replication origin binding protein, storage protein) compared to Olink 3K. Transketolase was identified in all SomaScan and MS-based platforms, whereas phosphatase activator was identified in SomaScan, MS-HAP Depletion and MS-Nanoparticle platforms.

**Figure 4:**
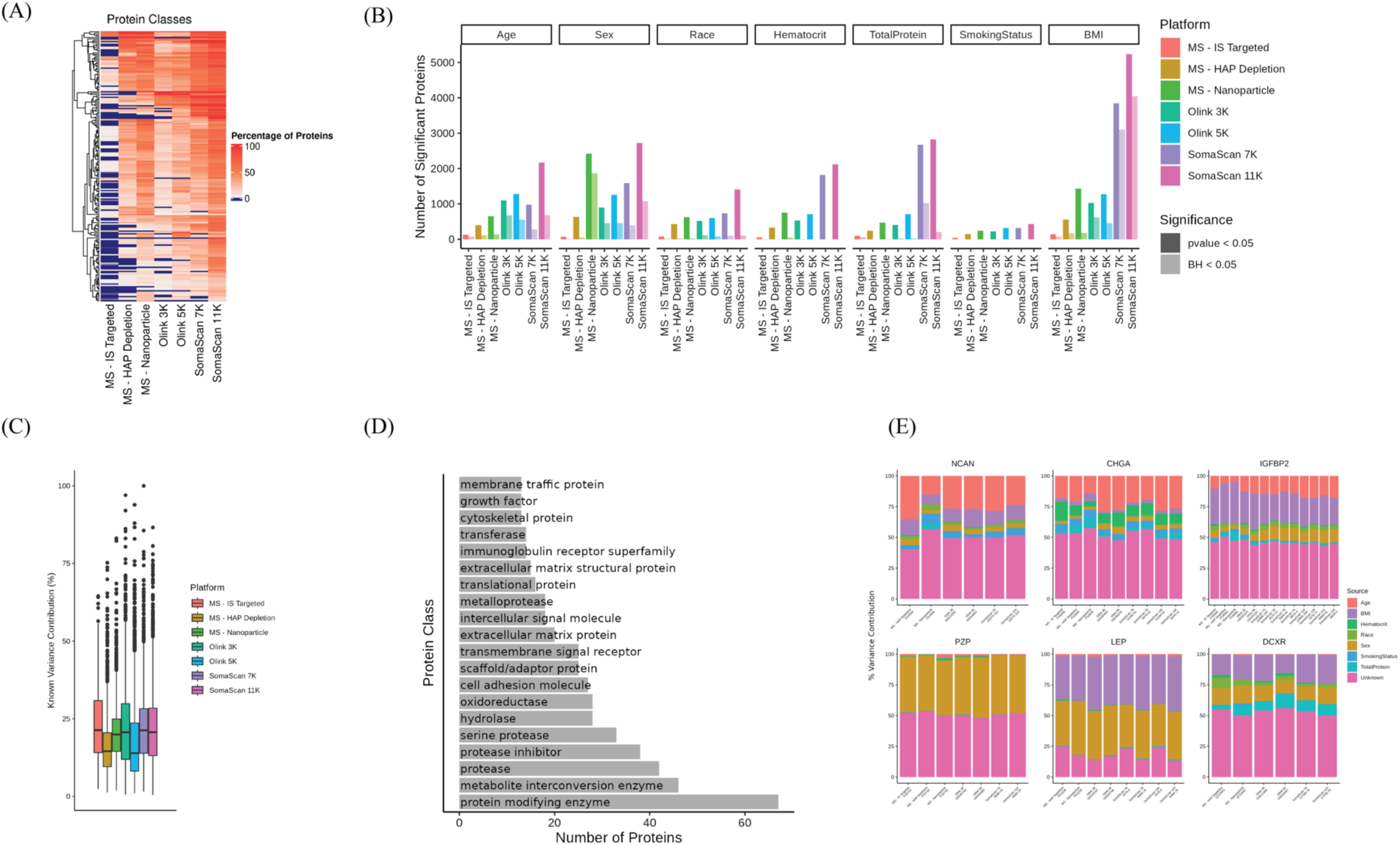
Platform protein categorizations, model results and variance analysis. (A) Percentage of proteins present in each PANTHER protein class across each of the platforms. (B) Number of identified protein markers across each of the platforms, for each predictor, based on a 0.05 p-value and 0.05 adjusted p-value cut-off. (C) Percent of variance unidentifiable from model parameters for each analyte mode not identifiable by included model parameters. (D) PANTHER protein classes are shown for proteins with greater than 50% unknown variance across all platforms except for MS-IS targeted. (E) Variance composition for proteins with greater than 60% known variance across all platforms except for MS-IS targeted.

To assess the biological relevance of these proteins, we employed a linear model that incorporated available metadata (age, sex, race, hematocrit, total protein, BMI, and smoking status) to identify significant markers associated with each factor across the various platforms. Figure 4B illustrates the number of significant markers identified by each platform, based on either p-values or Benjamini-Hochberg adjusted p-values (with p-val < 0.05 or p-adj < 0.05). SomaScan 11K, which covered the highest number of proteins and protein classes, identified the greatest number of biologically relevant markers compared to other platforms for age (2,170 and 685 p-val and p-adj, respectively), BMI (5,239 and 4,040 p-val and p-adj, respectively), and sex (2,726 and 1,074). MS-Nanoparticle also identified large number of sex related markers and majority of these markers were significant after the p-value adjustment procedure (2,427 and 1,873 p-val and p-adj, respectively). Supplementary Figure 7 highlights distribution of linear model coefficients categorized by their significance level. After SomaScan platforms, Olink 3K and 5K merged as the next most comprehensive assays for identifying markers related to age, sex, and BMI.

Our results show that the number of identified markers varied across platforms, regardless of the number of proteins they covered. Notably, the number of significant markers fluctuated between platforms depending on whether p-values or adjusted p-values were used. The distribution of p-values had a significant impact on the results after adjustment, with many platforms showing a notable reduction in the number of identified markers. For example, while the number of sex-related markers varied between Olink 3K, Olink 5K and SomaScan 7K at the p-value level, with SomaScan 7K identifying the highest number of markers, the number of markers became more comparable across these platforms after p-value adjustment. This reduction was most pronounced in MS-HAP Depletion platforms, where the number of corrected markers was substantially smaller, with 52 (p-adj), compared to the 632 (p-val) markers. Supplementary Figure 8 summarizes number of common and unique biological markers for each platform.

A variance decomposition analysis (Supplementary Figure 9) revealed that each platform captured unique biological factors and to varying degrees. By accounting for known factors of age, sex, race, hematocrit levels, total protein BMI and smoking status in our analysis, we evaluated the percentage of unexplained variance specific to each platform, as shown in Figure 4C. A substantial portion of the variance remained unexplained, suggesting the influence of additional biological factors beyond those included in our model. Two main factors of disease status and genomics association that have been reported^9^ to be highly important in explaining variance of plasma proteins are absent in our dataset as our cohort consists of healthy individuals and lacks genomics associations due to its small number of subjects. Figure 4D lists top 20 protein classes that are representative of proteins with more than 50% unexplained variance shared among all platforms. In our cohort with information on biological factors available, MS-IS Targeted, Olink 3K, MS-Nanoparticle, SomaScan 7K and 11K explained about the same amount of variance (19.9-21.3%) which is higher than Olink 5K, and MS-HAP depletion platforms (13.8-14.5%). Notably, despite covering the fewest proteins, MS - Targeted explained a significant amount of variance.

Figure 4E highlights examples of proteins, all exhibiting more than 40% explained variance contribution in all platforms. These examples demonstrate similar variance contribution of different platforms for candidate biomarkers of sex, age and BMI. For instance, Leptin (P41159), a circulating adipokine involved in regulating appetite, food intake, and fat distribution,^32^ is known to be influenced by sex hormones. Sex-related differences in leptin levels are well-documented, with women generally having higher concentrations than men.^33^ In obese individuals, leptin levels are elevated and correlate with BMI and body fat percentage.^34^ This aligns with our data, where biological factors such as sex and BMI explained 38.4% and 40.5% (in average among all platforms) of the variance associated with this protein across all platforms. Another example of sex biomarker is Pregnancy Zone Protein (P20742), where the sex factor alone explained up to 46.3% of the variance in PZP levels. This is consistent with strong sex-related differences in PZP plasma levels, with females having significantly higher levels than males.^11^ Chromogranin-A (P10645) and Insulin-like growth factor-binding protein 2 (P18065), two known markers of age,^35^ explain age variable with 22.9% and 13.3% or higher variance in all platforms. As expected IGFBP2, as insulin regulatory protein explains BMI the most^36^ and again this observation is consistent between all platforms. Similar patterns of variance decomposition were also observed across platforms for Neurocan core proteins (O14594) a predictive marker of age^37^ and L-xylulose reductase (Q7Z4W1) which participates in glucose metabolism.

### Age Related Markers

Among all biological factors analyzed in the study, we closely examined age related markers. Figure 5A represents the intersection of significant (p-adj < 0.05) age markers identified across all platforms. Olink 3K identified the highest number of age markers (669), followed by SomaScan 11K (628). Among the MS-based platforms, MS-Nanoparticle had the most age-related markers. In terms of the exclusive age markers identified by a platform, SomaScan 11K identified the greatest number of 282 markers that were not found by any other platform, followed by Olink 3K and Olink 5K, which identified 176 and 99 exclusive markers respectively. Seven proteins (P07998, P10645, P17936, P18065, P49747, Q15113, and Q9NQ79) appeared in all platforms including MS-IS Targeted. P18065 (IGFBP2) and P17936 (IGFBP3), insulin-like growth factor (IGF) binding proteins, are well-known markers of aging and have been identified in plasma proteomics studies as associated with aging. Both proteins are linked to cellular senescence and are likely involved in age-related physiological and pathological processes. P10645 (CHGA), P49747 (COMP) and Q9NQ79 (CKTAC1) markers of extreme agers has been recently reported as an age-associated protein.^35^ Q15113 has been also reported as aging protein discovered via quantitative mass spectrometry.^38^ P07998, is among Top 20 most significant SOMAmer reagents associated with chronological age.^39^

**Figure 5:**
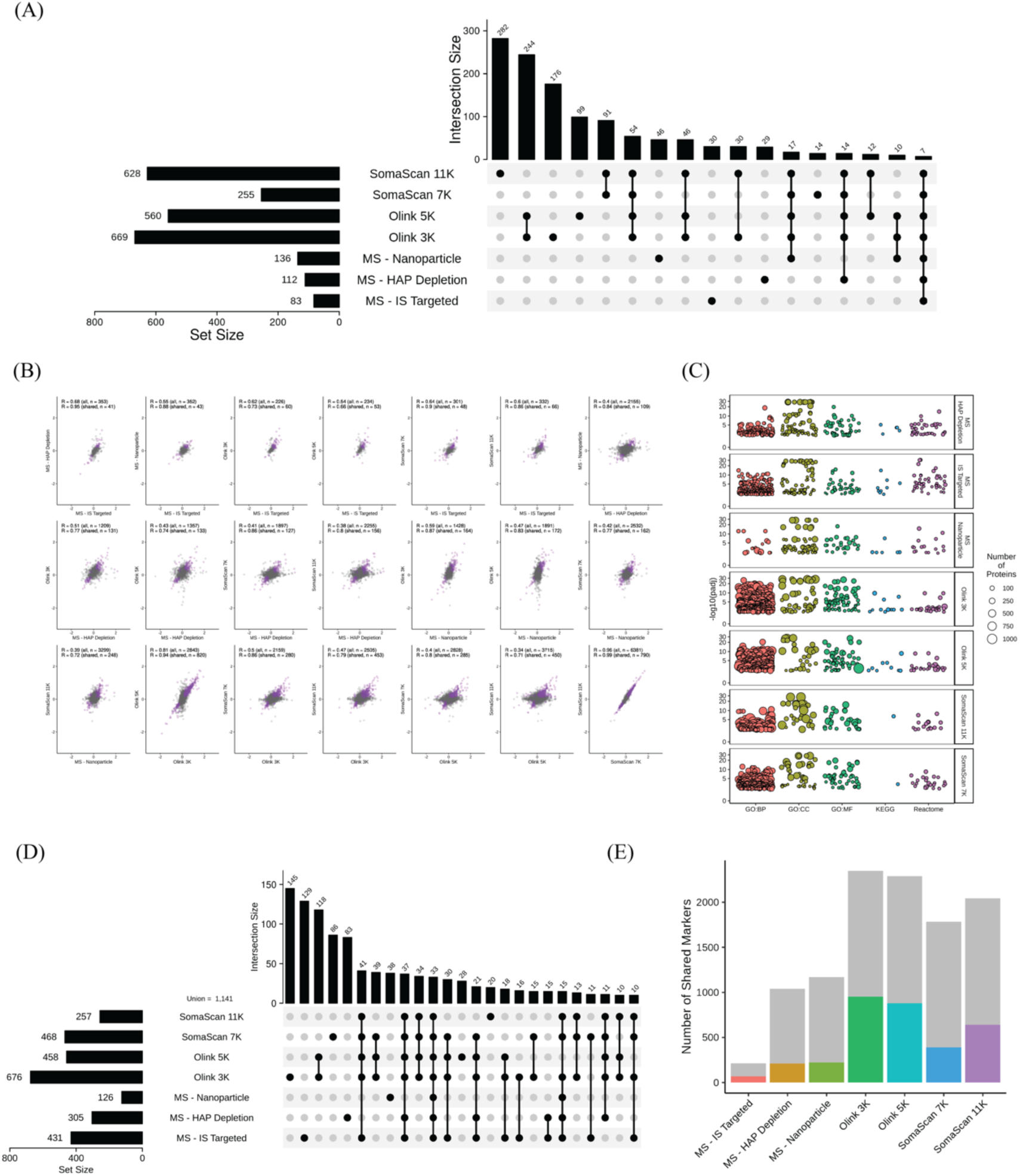
Age-related protein analysis and pathway enrichment. (A) UpSet plot showing the intersection of significant age markers across each platform. Significance determined by p-adjust < 0.05. (B) Correlation of coefficients between models for the shared aging marker proteins across different platforms. (C) Results of pathway enrichment analysis using significant age markers from each platform and union of proteins across all platforms as a background. GO, KEGG and Reactome pathway analysis of significant age markers from each of the platforms. Each bubble represents a term, and the size of the bubble indicates the number of involved markers. Only terms with significant p.adust < 0.01 are shown. (D) UpSet plot showing the intersection of significant age-related pathways identified across platforms. Significance determined by p-adjust < 0.01. (E) The number of UKB markers detected in both the platform of interest and the UKB dataset is shown in grey. The colored bars in the foreground indicate the number of UKB markers identifiable in both the platform of interest and the UKB dataset that were also identified as significant by our models.

Although correlation between shared proteins in platforms are quite low, correlation analysis between significant age markers common to platforms revealed a high degree of agreement (Figure 5B) with correlation of coefficients from 0.66 to 0.95 indicating a strong correlation among these markers. In contrast, the correlation between the coefficient of all shared proteins is less varied (between 0.34 to 0.68). This suggests that when a marker is deemed significant, it is likely to exhibit a strong correlation across multiple platforms.

Figure 5C shows pathway analysis of significant age markers, revealing distinct patterns across the various platforms. SomaScan 11K, SomaScan 7K and Olink 5K showed enrichment in GO terms related to molecular function, cellular component and biological process, along with Reactome pathways. In contrast, Olink 3K was primarily enriched in GO terms related to cellular components. The MS based platforms, MS-Nanoparticle and MS-HAP Depletion, exhibited enrichment in molecular function, cellular component, and Reactome pathways, but to a lesser extent than the affinity - based platforms. Overall, pathways enriched based on age related markers were substantially higher for affinity platforms.

We specifically studied how the most recent changes in the SomaScan and Olink technologies affected the number of age-related markers and their enriched pathways. In both the SomaScan and Olink technologies, we found the recent advancements in both technologies led to discrepancies between versions of the same technologies. For example, proteins that while detected in both were only significant in one of the versions of the platform and not the other.

We found that enrichment of age-related markers in Olink 3K was highly dependent on the background used. When using the Olink 3K universe as a background, we identified 12 enriched terms. However, when using the combined universes of Olink 3K and 5K as a background, we identified 491 additional terms to be enriched. We then investigated how using the union of detected proteins across all platforms as a background affected the enrichments. Using this method, we identified 426, 305, 127, 677, 459, 456, and 263 enriched terms in MS-IS Targeted, MS-HAP Depletion, MS-Nanoparticle, Olink 3K, Olink 5K, SomaScan 7K and SomaScan 11K respectively at the p-adjust < 0.01 level (Figure 5C), of which 127, 85, 39, 147, 29, 79, and 21 were unique to each platform. Thirty-three of these terms were commonly enriched across all platforms (Figure 5E). Most of these enriched terms were extracellular, binding, and adhesion related, however, IGF transport regulation, complement and coagulation cascades, and post translational protein phosphorylation were also highly conserved.

To assess the relevance of the age markers identified in this study, we compared the significant markers across all platforms with those reported in the UK Biobank Pharma Proteomics Project (UKB-PPP). The UKB-PPP dataset contains plasma proteomic profiles of 54,219 participants using the Olink Explore 3072 platform.^40^ This dataset is widely used in plasma proteomics research. Figure 5D displays the number of markers identified by each platform that were also present in the UKB dataset. Despite the limited size of the cohort included in our analysis, we observed overlap between the markers identified by each platform and those in the UKB dataset, reinforcing the biological relevance and reliability of the age - related markers identified across the different platforms.

## Discussion

Recent advancements in protein profiling technologies have significantly enhanced our ability to understand the plasma proteome. Affinity proteomics has seen a remarkable increase in the number of available assays. For instance, the proteome coverage of SomaScan has expanded from approximately 800 assays in 2009 to 11,000 assays as of beginning of 2025. Similarly, Olink has increased the number of assays to over 5,400 with the introduction of Olink Explore HT - a substantial increase from fewer than 1,000 assays available approximately 15 years ago.

Concurrently, mass spectrometry has made notable progress, improving sample throughput, quantification, and increasing the depth of coverage in plasma proteome. These advancements have facilitated the development of robust, automated, and scalable workflows for comprehensive plasma proteome profiling. One such workflow employed in this study utilizes HAP depletion and deep coverage ion mobility data-independent acquisition (DIA) methods.^16^ Additionally, nanoparticle-based enrichment techniques have been integrated into mass spectrometry workflows for plasma proteomics. A notable example is the Seer Proteograph™ XT, introduced in 2020, which uses magnetic nanoparticles with different surface chemistries to selectively enrich proteins based on their physicochemical properties prior to LC-MS/MS analysis.^20^ This approach has markedly enhanced the coverage of the plasma soluble proteome.^20,41,42^

Other recent methodologies of depletion or enrichment that have been effective in increasing plasma proteome coverage include perCA depletion,^18^ Enrich and Enrichplus kits containing enrichment beads from Preomics, and the proprietary P2 Plasma Enrichment method from Biognosys, Collectively, these innovations have greatly expanded the depth of proteome analysis, enabling the identification and quantification of previously elusive low-abundance proteins in plasma.

As interest in molecular profiling of plasma for biomarker discovery and target identification grows, both affinity-based and mass spectrometry platforms have become essential tools in research and clinical applications. While several previous studies have compared two or three proteomics platforms across healthy and diseased cohorts, our study focuses on a well-defined healthy cohort with high level of comprehensiveness by evaluating seven proteomic technologies on a single set of plasma samples (Figure 1). The seven platforms assessed here represent five different proteomic approaches and collectively encompass an extensive array of over 29,013 assays and 16,493 unique proteins. While affinity-based assays were able to detect very low-abundance proteins, their discovery panels used in our study provide only relative quantification rather than absolute concentrations. Absolute quantification can be achieved by using methods incorporating standard curves or quantitative internal standards, such as the MS-IS Targeted method used in this study. While it identified fewer proteins, it uniquely provided absolute quantification values which further facilitates downstream workflow for identified markers or targets. We believe that our selection of four affinity platforms, two discovery MS platforms and one targeted MS platform, brings a comprehensive view of current proteomics platforms, with the absolute quantification offered by targeted MS serving as a valuable reference. List of all identified proteins from seven platforms and their presence in published datasets of FDA, UK Biobank and HPA are listed in Supplementary Table S6.

The optimal application of plasma proteomics for biomarker and target discovery and validation requires a deep understanding of both the platform’s coverage and its technical characteristics. SomaScan 11K offers the most comprehensive coverage of the plasma proteome, making it the best platform available for discovery studies with the objective of identifying previously unknown biological roles for the largest set of proteins. This largest coverage is even more noteworthy in view of the practically complete detectability of healthy plasma levels of the covered proteins in the linear range of the assay. While mass spectrometry now provides greater coverage than before, it still falls short in robust detection of low-abundance proteins due to its technological limitations. However, the high-resolution data now available on precursor ions offers new opportunities for the direct identification of isoforms and post-translational modifications (PTMs) without the need for prior enrichment.

A recent technical evaluation of the latest versions of the affinity proteomics platforms used for plasma analysis highlights key differences in their performance, especially concerning the technical coefficients of variation (CV) and completeness of data.^28^ The primary focus of our evaluation is the gap between the technical CVs and the total CVs observed in the cohort, as this gap can reveal the extent to which technical variability interferes with detecting biological changes of interest (Figure 2B). This gap is particularly favourable for platforms like SomaScan 7K and 11K, Olink 3K, and MS-IS targeted assays. In general, affinity-based platforms such as SomaScan and Olink tend to exhibit lower technical CVs compared to discovery mass spectrometry (MS), indicating superior precision. However, the Olink 5K platform demonstrates a technical CV comparable to that of discovery MS, with a coefficient approximately double that of the other platforms.

Another significant observation from our analysis is that Olink 5K assays with fewer than one-third missing values exhibit median CV about half of full-assay platform. However, filtering based on eLOD removes more than half of the assays in the platform, representing about 2,000 proteins. The analysis also found that data completeness, as determined by the number of missing values or values below the estimated limit of detection (eLOD), is inversely related to the technical CV across all platforms. This suggests that assays with higher rates of missing data, which are often linked to low-abundance proteins, tend to exhibit higher variability, further complicating the interpretation of results.

The comprehensive coverage of SomaScan 11K is particularly notable in its ability to capture the highest number of FDA-approved protein biomarkers identified to date (Figure 2C). As a result, SomaScan provides valuable opportunities for deeper insights into the biology of plasma samples. Our findings further support this advantage, as SomaScan demonstrated the ability to identify a greater number of significant biological markers, offering a more nuanced understanding of the underlying biological processes. Untargeted mass spectrometry, particularly when combined with complexity reduction strategies such as HAP (high-abundance protein) depletion and the use of nanoparticles, has proven effective in addressing the challenges of identifying low-abundance and clinically relevant proteins.^16,42^ These strategies enable a more robust identification of these proteins, which are often crucial for understanding disease mechanisms. In fact, a significant percentage (72.8%) of FDA-approved biomarkers were identified across the MS platforms used in this study (Figure 2C), further supporting their extended coverage and ability to capture a broader spectrum of biologically relevant targets. This highlights the potential of untargeted MS, when optimized with advanced techniques, to provide valuable insights into complex proteomic landscapes.

With numerous publications comparing different proteomics platforms,^31,25,28,43,44,45^ it is no longer surprising to see low levels of correlation between these technologies. The lack of correlation is understandable, though it is difficult to pinpoint specific causes for specific situations as multiple factors can influence protein identification. Affinity-based platforms identify proteins based on three-dimensional binding epitopes that can be different from assay to assay even when the binding probes were raised against the same target protein construct. Bottom-up mass spectrometry (whether targeted or untargeted) relies on the detection of proteotypic peptides of denatured and digested proteins; these again can be different from one MS assay to the other. Based on the various binding and sequence motifs detected by the different proteomic assays, the protein and proteoform specificity profiles of the platforms can be different, which in turn, can explain diminished correlation.

Beyond the above fundamental methodological differences, the precision of the platform can also contribute to the observed variability in correlations. We found that proteins with higher technical CVs (>20%) tend to show lower correlations with other platforms, while those with lower CVs (<20%) exhibit relatively higher correlations (Figure 3C). This confirms that the precision of the platform can, to some degree, explain the differences in correlation, but it is not the sole factor. The separation of the CV-dependent correlation patterns is particularly pronounced for some platforms, such as SomaScan 7K and 11K, but less evident for the MS-based platform that uses nanoparticle-based complexity reduction. This illustrates how differences in identification approach and quantification precision of a platform can result in a wide range of correlation and agreeability with other platforms. Previous studies have shown that median Spearman’s correlations for overlapping proteins between SomaScan and Olink ranges from 0.20 to 0.44. In contrast, direct comparisons between affinity-based and MS-based platforms are more limited. To our knowledge, one prior study has directly compared SomaScan, Olink, and MS-based platforms using plasma samples.^46^ This study found a median Pearson correlation of 0.55 between MS and Olink, and 0.59 between MS and SomaScan. Notably, the number of proteins analyzed in each correlation was 295 for MS - Olink and 952 for MS - SomaScan, reflecting the use of earlier versions of the platforms in that study. Additionally, the MS platform employed TMT - based DDA analysis, which differs from the DIA and targeted approaches used in our study. This underscores the need for more comparative studies between mass spectrometry and affinity - based platforms, particularly with the emergence of new mass spectrometry technologies.

Intra-platform correlations between different assays for the same protein, such as assays using different SOMAmers developed for the same target in the Somacan platform, can often be low. Our data shows that the distribution of the correlation coefficient is bimodal with assays for one group of targets showing strong correlations (rho > 0.7) but for the other group assays correlating weakly (-0.1 < rho < 0.7). This phenomenon can be attributed to the fact that different SOMAmers may have different binding profiles across distinct proteoforms of the same protein, especially since SomaScan includes aptamers selected against different expression constructs of the same protein. As a result, different SomaScan assays targeting the same protein may preferentially measure different isoforms or variants of the protein, leading to discrepancies in the measured levels. It is important to note, however, that these discrepancies are also sample dependent: they will manifest only in samples that have different proteoform profiles. While non-agreement of different assays targeting the same protein sounds like a limitation of a proteomics platform, it also reveals valuable information by pointing to potential proteoform heterogeneity between the measured samples.

To investigate protein or proteoform selectivity discrepancies, pull-down experiments can be useful to isolate the plasma proteins bound by the affinity probes, followed by the identification and quantitation of the captured proteins. Published efforts using mass spectrometry have so far demonstrated the enrichment of the intended protein targets for a subset of SOMAmers^47^ but full protein selectivity characterization of entire affinity proteomics platforms is still elusive. Proteoform selectivity determinations are even more challenging and would likely not be widely achievable until protein analytical technologies (e.g., top-down MS) evolve to the level of routine proteoform characterization from very small sample sizes.

In addition to the technical evaluations of the platforms, there are notable differences in their biological assessments. Each platform provides unique insights into biological factors, as demonstrated by the number of significant proteins identified for each factor (Figure 4B) and the variance they account for (Figure 4C). While many protein classes are shared across platforms, the number of proteins within each class can either enhance or limit the capacity to elucidate biological factors to different extents (Figure 4A). This highlights the importance of selecting the appropriate platform based on the specific biological questions at hand.

Change in number of significant protein markers behave differently after p-value adjustment across different platforms. For age-related markers, both MS-Nanoparticle and MS-HAP platforms initially present hundreds of markers with p-values less than 0.05. However, only a fraction of these markers— specifically, 136 and 112 markers—remain significant after p-value adjustment. Interestingly, this number of adjusted significant markers is comparable to the number of significant markers identified by MS-targeted analysis, despite the fact that only about 500 proteins were identified in the targeted MS analysis. With increase in number of proteins identified in proteomics platforms, the stringency of statistical corrections may have a considerable impact on the interpretation of proteomics data across platforms.

The variance explained in our study also varies across different platforms, though the differences are relatively minor. Overall, the proportion of variance explained by biological factors in our study is lower than what is typically explained by proteomics in plasma. Other studies that incorporated genomic data from the cohort, as well as information on disease status, were able to explain a larger portion of the plasma variance through genome associations and disease factors. However, in our study, age, sex, and BMI emerged as the primary contributors to the variance observed in plasma proteomics. For shared markers across all platforms, a similar pattern of variance decomposition was observed, indicating that these biological factors consistently play a dominant role in shaping the plasma proteome across different proteomics technologies.

Plasma aging markers have been extensively studied, but our research provides a comprehensive overview of aging-related proteins in plasma across seven different platforms for the first time. Our findings reveal that newer platforms, such as SomaScan 11K and Olink 5K, introduce a broader array of age-related markers compared to their predecessors. Specifically, SomaScan 11K identifies 279 markers that were not present in SomaScan 7K, while Olink 5K detects 98 unique proteins not found in previous versions. Overall, affinity-based platforms contributed the largest number of both unique and shared significant aging markers. In contrast, discovery and targeted mass spectrometry (MS) platforms identified fewer significant markers, but each added valuable proteins that were not covered by the affinity proteomics platforms (Supplementary Figure 10).

While correlations between platforms are generally weak, pairwise comparisons of shared markers demonstrate strong agreement in the coefficients of significant markers (p-value < 0.05). This suggests a higher level of concordance across platforms when identifying key age-related markers. Additionally, when cross-referencing our data with the recent UK Biobank proteomics analysis, we observe platform-specific biases in marker identification. Notably, the Olink 3K and 5K platforms exhibit the highest overlap with age-related markers identified in the UK Biobank, as Olink was the platform used for their proteomic measurements.

This study presents a comprehensive list of aging-related proteins in plasma. When cross-referencing our identified markers with those reported in the literature, we find substantial overlap, suggesting that our study has successfully captured many of the previously discovered markers, regardless of the platform used for protein profiling. Specifically, 78% of the proteins from the recently published aging clock based on the UK Biobank and the Olink platform^48^ were also identified in our study, including all but one of their top 20 ProtAge signatures. Similarly, for age-related markers identified using the SomaScan platform, over 92% of the markers shared across four large cohorts measured with SomaScan^49^ were also detected in our dataset, with 94% of them overlapping with centenarian markers. Furthermore, 80% of the centenarian markers discovered via MS^35^ were also present in our findings.

The number of significant pathways associated with aging markers varies considerably across platforms, likely due to differences in the number of markers identified (p < 0.05). For instance, the SomaScan 11K panel, with over 2,000 markers, identifies more pathways compared to MS-based targeted approaches, which typically have fewer markers (e.g., 133 in the IS Targeted platform). Additionally, the nature of the proteins detected influences downstream pathway analysis and enrichment. Targeted platforms, such as SomaScan, often select panels based on known biological pathways, leading to the identification of established pathways. In contrast, untargeted MS approaches may yield a similar number of significant markers but may not highlight known pathways. However, these untargeted methods have the advantage of enabling the discovery of novel biomarkers that fall outside of predefined pathways.

In conclusion, this study provides a comprehensive comparison of different proteomic platforms, highlighting the strengths and weaknesses of each in characterizing the plasma proteome. Ultimately, the choice of platform depends on the specific research question and the desired depth of proteomic analysis. As proteomic technologies continue to evolve, ongoing comparative studies will be crucial for refining our understanding of their capabilities and limitations in plasma proteome profiling. These insights are vital for improving biomarker discovery, advancing clinical applications, and deepening our understanding of complex biological processes. By continuing to evaluate and compare these evolving technologies, researchers can select the most appropriate tools for their investigations and unlock the full potential of plasma proteomics.

## Methods

### Ethics

Informed consent was obtained from all subjects.

### Study Enrolment

Subjects were selected from among regular plasma donors of Access Biologicals. 80 subjects (40 female and 40 male) were selected in two groups of young (18-22 years old) and aged (55-65 years old). Subjects were considered healthy since they were from regular plasma donor participants. Information on their medications, medical condition, fasting time, smoking status as well as biometric data (BMI), blood pressure, temperature, hematocrit level, total protein, age, gender and race were collected.

### Plasma Collection

Plasma was collected through plasmapheresis with Sodium citrate as anticoagulant from each participant.

### Plasma profiling - SomaLogic

Plasma samples were analyzed in two versions of the SomaScan proteomics platform, SomaScan v4.1 (7K) and v5.0 (11K), at Somalogic (Boulder, CO, USA). The two platform versions share the same assay technology but differ in the number of assays included (7,288 vs 10,776 human protein assays, respectively).

The technology is based on Somalogic’s proprietary Slow Off-rate Modified Aptamers (SOMAmers) that bind to structural epitopes on proteins. SOMAmers include a fluorophore and a photocleavable biotin moiety in addition to the binding probe single-stranded DNA sequence that contains modified bases to promote protein binding. During the assay, the SOMAmer reagents are pre-immobilized onto streptavidin beads and used to capture target proteins from biological samples. Unbound proteins are washed away, and captured proteins are biotinylated using NHS-biotin. UV light is used to cleave the photosensiitive linker, releasing complexes back into solution in the presence of a high concentration of universal polyanionic competitor. Complexes and some free proteins that dissociated are captured onto new streptavidin beads. After washing, SOMAmer reagents are eluted from the beads by denaturing the proteins. The eluate is placed onto a custom Agilent microarray with probes complimentary to each SOMAmer reagent for overnight hybridization. Slides are washed and read in an Agilent microarray scanner. The resulting RFU values reflect the amount of target epitope in the initial samples.

SomaScan Assay data are normalized using hybridization controls followed by median signal normalization across calibrator replicates within the run. Plate scale factors and calibration scale factors based on the calibrator replicates and external reference values are used to adjust for overall signal intensity differences between runs and SOMAmer reagent-specific assay differences, respectively. Median signal normalization is performed using Adaptive Normalization by Maximum Likelihood (ANML).

### Protein Profiling - Olink

Plasma samples were analyzed in two versions of the Olink proteomics platform, Olink® Explore 3072 and Explore HT (Olink Proteomics AB, Uppsala, Sweden) at Olink Analysis Services in Boston (MA, USA). The two platform versions share the same assay technology but differ in the number of assays (3,072 vs 5,416, respectively) as well as sample requirement and throughput.

The shared underlying technology is based on Proximity Extension Assay (PEA) (Assarsson et al, 2014), coupled with next-generation sequencing (NGS) as readout. In brief, pairs of oligonucleotide-labeled antibody probes against the same protein bind to their target, bringing the complementary oligonucleotides in close proximity and allowing for their hybridization. The addition of a DNA polymerase leads to the extension of the hybridized oligonucleotides, generating a unique protein identification “barcode”. Next, library preparation adds sample identification indexes and the required nucleotides for Illumina sequencing.

Prior to sequencing using the Illumina® NovaSeq™ 6000/NextSeq™ 550/NextSeq™ 2000, libraries go through a bead-based purification step and the quality is assessed using the Agilent 2100 Bioanalyzer (Agilent Technologies, Palo Alto, CA). The raw output data is quality controlled, normalized and converted into Normalized Protein expression (NPX) values, Olink’s proprietary unit of relative abundance. Three internal controls are spiked into every sample and are used to monitor the performance of the three main steps in the protocol: an incubation control, an extension control and an amplification control. In parallel with the samples, the protocol is performed on a set of external controls: Two sample controls, three negative controls and three plate controls (PCs).

The PCs are used for between-plate normalization. Quality control is performed for each sample plate on both the samples (using the spiked internal controls) and the external controls. All assay validation data (detection limits, intra- and inter-assay precision data, predefined values, etc.) are available on manufacturer’s website (www.olink.com).

### Protein Profiling - Biognosys TrueDiscovery™ Pipeline

All solvents were HPLC-grade from Sigma-Aldrich and all chemicals where not stated otherwise were obtained from Sigma-Aldrich. Plasma samples were shipped frozen by Alkahest. Samples were depleted using a Multiple Affinity Removal Column Human 14 (Agilent) column. Samples were prepared for LC-MS/MS according to Biognosys’ SOP which includes reduction, alkylation and digestion to peptides using trypsin (Promega, 1:50 protease to total protein ratio) per sample overnight at 37 °C. Peptides were desalted using a C18 HLB µElution plate (Waters) according to the manufacturer’s instructions and dried down using a SpeedVac system. Peptides were resuspended in 1 % acetonitrile and 0.1 % formic acid and spiked with Biognosys’ iRTkit calibration peptides. Peptide concentrations were determined using a UV/VIS Spectrometer at 280 nm (SPECTROstarNano, BMG Labtech).

For DIA LC-MS/MS measurements, 3.5 µg of peptides per sample were injected to an in-house packed reversed phase column on a ThermoScientific™EASY-nLC™1200 nano-liquid chromatography system connected to a ThermoScientific™Orbitrap™Exploris480™mass spectrometer equipped with a NanosprayFlex™ ion source and a FAIMS Pro™ ion mobility device (ThermoScientific™). LC solvents were A: water with 0.1 % FA; B: 80 % acetonitrile, 0.1 % FA in water. The nonlinear LC gradient was 1 – 50 % solvent B in 210 minutes followed by a column washing step in 90 % B for 10 minutes, and a final equilibration step of 1 % B for 8 minutes at 60 °C with a flow rate set to 250 nL/min. The FAIMS-DIA method consisted per applied compensation voltage of one full range MS1 scan and 34 DIA segments as DIA method consisted per applied compensation voltage of one full range MS1 scan and 34 DIA segments as adopted from Bruderer et al. [A] & Tognetti et al. [B].

A directDIA™ spectral library was generated by searching the HRM mass spectrometric data using Spectronaut (Biognosys, version 16.2), the false discovery rate on peptide and protein level was set to 1 %. A human UniProt. Fasta database (Homo sapiens, 2022-07-01) was used for the search engine, allowing for 2 missed cleavages and variable modifications (N-term acetylation, methionine oxidation, deamidation (NQ) and ammonia-loss). The results were combined with a proprietary deep spectral library for human plasma using Spectronaut.

Raw mass spectrometric data were first converted using the HTRMS Converter (version 15.6, Biognosys) and then analyzed using Spectronaut (Biognosys, version 16.2) with the default settings, but Qvalue filtering with background signal as imputation method was enabled and the hybrid spectral library generated in this study was used. Default settings included peptide and protein level false discovery rate control at 1% and cross-run normalization using global normalization on the median.

For testing of differential protein abundance, protein intensities for each protein were analyzed using a two-sample Student’s t-test. P-values were corrected for overall FDR using the q-value approach. The following thresholds were applied for candidate identification: q-value < 0.05; absolute average log2 ratio > 0.58 (fold-change > 1.5). Distance in heat maps was calculated using the “manhattan” method, the clustering using “ward.D” for both axes. Principal component analysis was conducted in R using prcompand a modified ggbiplotfunction for plotting, and partial least squares discriminant analysis was performed using mixOMICS package. Functional analysis was performed using String-db(string-db.org, version 11.5). Topology of candidate proteins was visualized using Protter. General plotting was done in R using ggplot2package.

### Protein Profiling - Seer Proteograph

Samples were processed with the Proteograph XT Assay [PMID: 32699280, PMID: 35275789]. In brief, 240 µL from each sample was transferred to Seer Sample Tubes for processing with the Proteograph XT Assay kit. Plasma proteins were quantitatively captured in nanoparticle (NP) associated protein coronas. Proteins were subsequently denatured, reduced, alkylated and subjected to proteolytic digestion (trypsin and LysC). Peptides were purified and yields were determined (Thermo Fisher Scientific catalog #23290). Peptides were dried down overnight with a vacuum concentrator and reconstituted with a reconstitution buffer to a concentration of 50 ng/µL.

For Data-Independent Acquisition (DIA), 8 µL of reconstituted peptide mixture from each NP preparation was analyzed resulting in a constant 400ng mass MS injection between NP A and NP B samples. Each sample was analyzed with a Vanquish NEO nanoLC system coupled with a Orbitrap TM Astral TM(Thermo Fisher, Germany) mass spectrometer using a trap-and-elute configuration. First, the peptides were loaded onto an AcclaimTM PepMapTM 100 C18 (0.3 mm ID x 5 mm) trap column and then separated on a 50 cm µPACTM analytical column (PharmaFluidics, Belgium) at a flow rate of 1 µL/min using a gradient of 5 – 25% solvent B (0.1% FA, 100 % ACN) mixed into solvent A (0.1% FA, 100% water) over 22 min, resulting in a 33 min total run time. The mass spectrometer was operated in DIA mode with MS1 scanning and MS2 precursor isolation windows between 380-980 m/z. MS1 scans were performed in the Orbitrap detector at 240,000 R every 0.6 seconds with a 5 ms ion injection time or 500% AGC (500,000 ion) target. Two-hundred fixed window MS2 DIA scans were collected at the Astral detector per cycle with 3 Th precursor isolation windows, 25% normalized collision energy, and 5 ms ion injection times with a 500% (50,000 ion) active gain control maximum. MS2 scans were collected from 150-2000 m/z.

Raw mass spectral files were processed using DIA-NN search engine, v1.8.1 using a Homo Sapiens FASTA file containing canonical reviewed and unreviewed proteins. Library-free search was performed In silico based on the input UniProt reference database listed above with Match Between Runs (MBR) enabled. A 1% FDR filtering for identification on peptide/protein group level and a quantification strategy of Robust LC (high precision) was used. DIA-NN search parameters included N-term M excision fixed modification, C carbamidomethylation fixed modification, minimum Peptide Length 7, maximum Peptide Length 30, minimum Precursor Charge 1, maximum Precursor Charge 4, minimum Precursor m/z 300, maximum Precursor m/z 1800, minimum Fragmentation Ion m/z 200, and maximum Fragment Ion m/z 1800. Precursor normalization was performed using calibration peptides (PepCal) spiked into each MS run to estimate MS drift. Panel Rollup was performed using MaxLFQ treating each nanoparticle as independent species per precursor via R package iq.^50^

Protein Profiling – SureQuant Undepleted human plasma was processed using the Thermo Scientific™ EasyPep™ MS Sample Prep Kit. A set of 804 SIL peptides from PQ500 (Biognosys PN# Ki-3019-96) was spiked at approximately 80 fmol (median value) into 1 µg of plasma tryptic digest. The volume corresponding to 1 µg of the digest was injected for LC-MS/MS analyses on an Orbitrap Exploris 480 mass spectrometer (Thermo Fisher Scientific) coupled with a Vanquish Neo UHPLC system (Thermo Fisher Scientific).

Chromatographic separations were performed using a 0.5 cm C18 PepMap™ Neo Trap Cartridge column (5 µm, 100 Å, 300 µm inner diameter; Thermo Fisher Scientific Cat# 174500) and a 15 cm C18 EASY-Spray™ HPLC column (2 µm, 100 Å, 150 µm inner diameter; Thermo Fisher Scientific Cat# ES906). Peptides were separated over a gradient from 2% to 31.5% 80% acetonitrile with 0.1% formic acid over 60 minutes.

To implement this method, the custom SureQuant acquisition template available in the Thermo Orbitrap Exploris was utilized. In the ’watch’ mode of the SureQuant method, the MS1 resolution was set to 120k to monitor the predefined optimal precursor ions of the internal standard (IS), which were included in the targeted mass filter. This was followed by the recognition of heavy peptide precursors and fragments from the list at a low resolution of 7.5k, with HCD collision energy set to 27% and a maximum injection time of 10 ms. The detection of the IS triggered the ’quant’ mode, which required at least five product ions to initiate an offset scan at a resolution of 60k, with HCD collision energy at 27%, a normalized AGC target of 1000%, and a maximum injection time of 116 ms in profile mode. Data analysis from SureQuant acquisitions was performed using SpectroDive™ (Biognosys).

### Dilution linearity

Two pooled plasma samples, one from young (18-22 year-old, pool of 20) and one from aged (55-65 year old, pool of 20) healthy males, were diluted 3x and 9x with PBS. At least three technical replicates of each of the undiluted samples, their two dilutions, and the buffer were tested by the proteomic platforms. Pearson correlation coefficient (r) of the resulting protein level versus sample relative concentration data was calculated for each measured protein and used to characterize the linearity of the platforms in the range of healthy plasma protein levels. Estimated Limit of Detection (eLOD) was calculated as mean + 3*SD of buffer controls.

### Data Analysis

Multi-UniProt Accession ID analytes were addressed by sorting the UniProt Accession IDs of these analytes in order to maintain consistency between platforms.

For the affinity-based platforms, eLOD values for each protein were calculated dependent on the platform. For SomaScan 7K and 11K platforms, eLOD values were calculated from buffer controls using 3.3*sd + mean. For Olink 3K, eLOD values were provided with the data. For Olink 5K, eLOD values were calculated using olink_lod function from the OlinkAnalyze 3.8.2 R package^51^ using the fixed LOD method.

Data completeness for each analyte was calculated across all platforms, defined by the percentage of analytes that were detected in each sample. For MS-based platforms, raw measurements were used. For affinity-based platforms, pre and post eLOD filtered values were used.

Technical and total CV percentage was calculated for each analyte across all platforms. For the SomaScan 7K assay dataset, technical CV was calculated using duplicates of eight samples. For the SomaScan 11K assay dataset, technical CV was calculated using four samples with four technical replicates each. This included one aged female bridging sample, one young female bridging sample, a bridging sample pool of 20 aged male subjects, and a bridging sample pool of 20 young male subjects. For Olink 3K, depleted Mass Spectrometry data, and SureQuant data, technical CV was calculated using three samples tested in triplicates. For Olink 5K and Seer, technical CV was calculated using two pools of samples, aged male and young male, each tested in six replicates. Total CV percentages were calculated for each analyte across all platforms, using 78 shared samples. Unique proteins were determined using unique UniProt Accession IDs. For platforms in which a single UniProt Accession ID was represented by multiple analytes, all analytes were kept separate in the analysis until UniProt Accession IDs were necessary to work with. To compare platforms to one another, we calculated correlation for analytes between platforms using spearman’s rank correlation. For instances in which there was a one-to-many or many-to-many relationship of analytes representing the same UniProt Accession IDs, correlations were calculated using all combinations of analytes. For SomaScan 7K and 11K, due to the analytes of 11K being a superset of the 7K analytes, we performed a one-to-one comparison of the same analytes Multivariate linear modeling was performed in R v4.3.2^52^ with lm according to the formula Log2(Protein Measurement) ∼ Age + Gender + Race + Hematocrit + TotalProtein + SmokingStatus + BMI. The protein measurement used was platform dependent. The mass spectrometry datasets were filtered for at least 2/3rds data completeness within each age group. The Olink datasets were left unfiltered. The SomaScan datasets were filtered based on eLOD values. Pathway enrichment was performed using topGO ^53^ v2.54.0, ReactomePA^54^ v1.46.0 and clusterProfiler^55,56^ v4.10.1 using proteins identified with a p-value less than 0.05. The total set of detected proteins for each platform was used as a background. Plots were generated using the R package ggplot2^57^ v3.5.1 Heatmaps and UpSet plots were generated using the R package ComplexHeatmap^58,59^ v2.18.0.

## Contributions

DK, AP, BS, SA analysis of data, SC, BS, SA writing the paper, BS sample management, BN data acquisition. All authors critically revised, edited, and approved the final manuscript.

---

**Figure S1:**
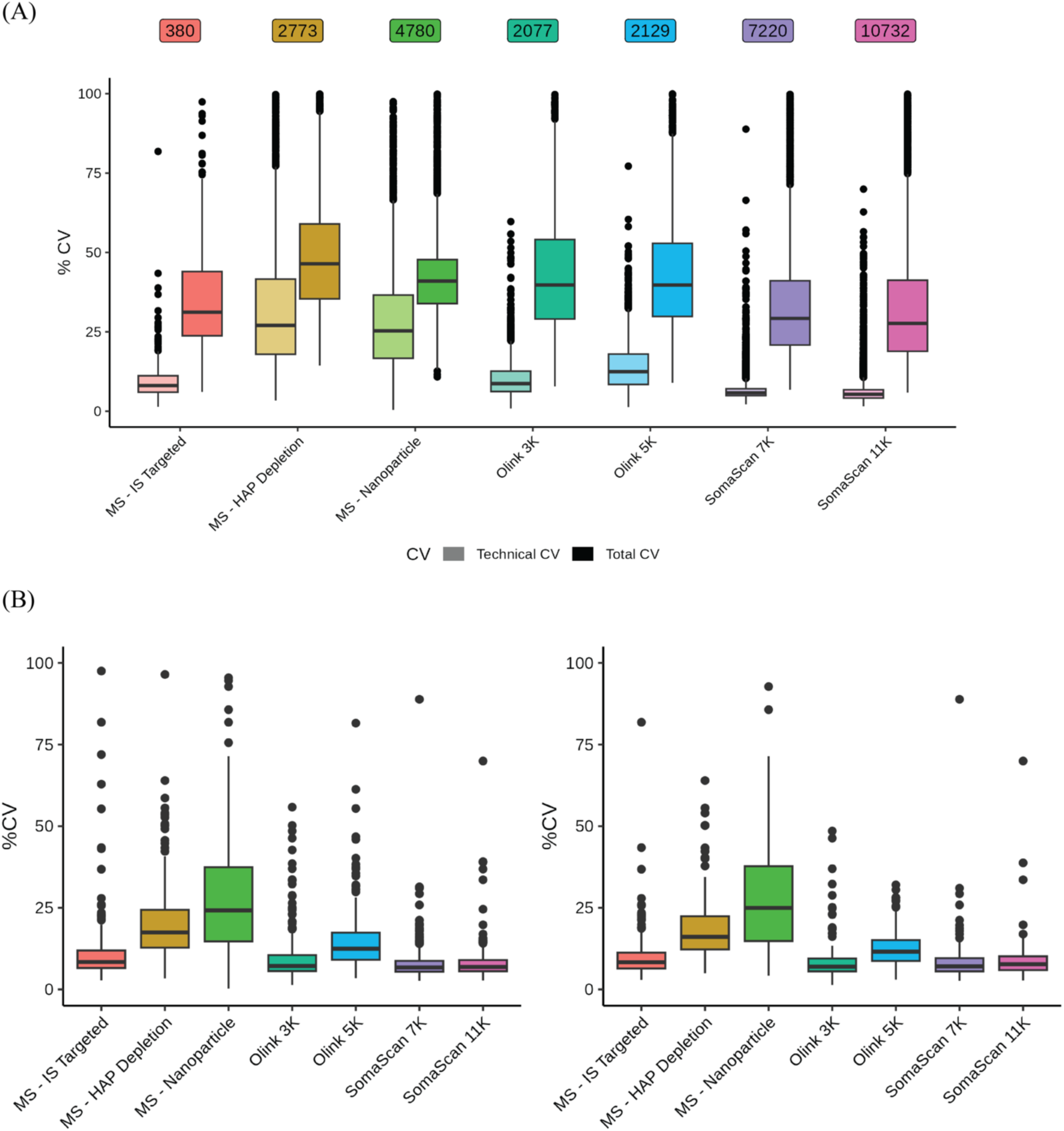
Influence of Filtering on CVs. (A) Technical and total CVs shown for each analyte on each platform, filtered for appropriate eLOD calculation and two-thirds data completeness. Technical CVs were calculated using technical replicates of n = 2 or n = 3, depending on the platform. Total CVs were calculated using the shared 78 subjects. (B) Technical CVs shown for each analyte corresponding to the 254 proteins detected in all platforms. The figure on the left displays unfiltered data, while the figure on the right corresponds to data filtered according to eLOD and two-thirds data completeness.

**Figure S2.**
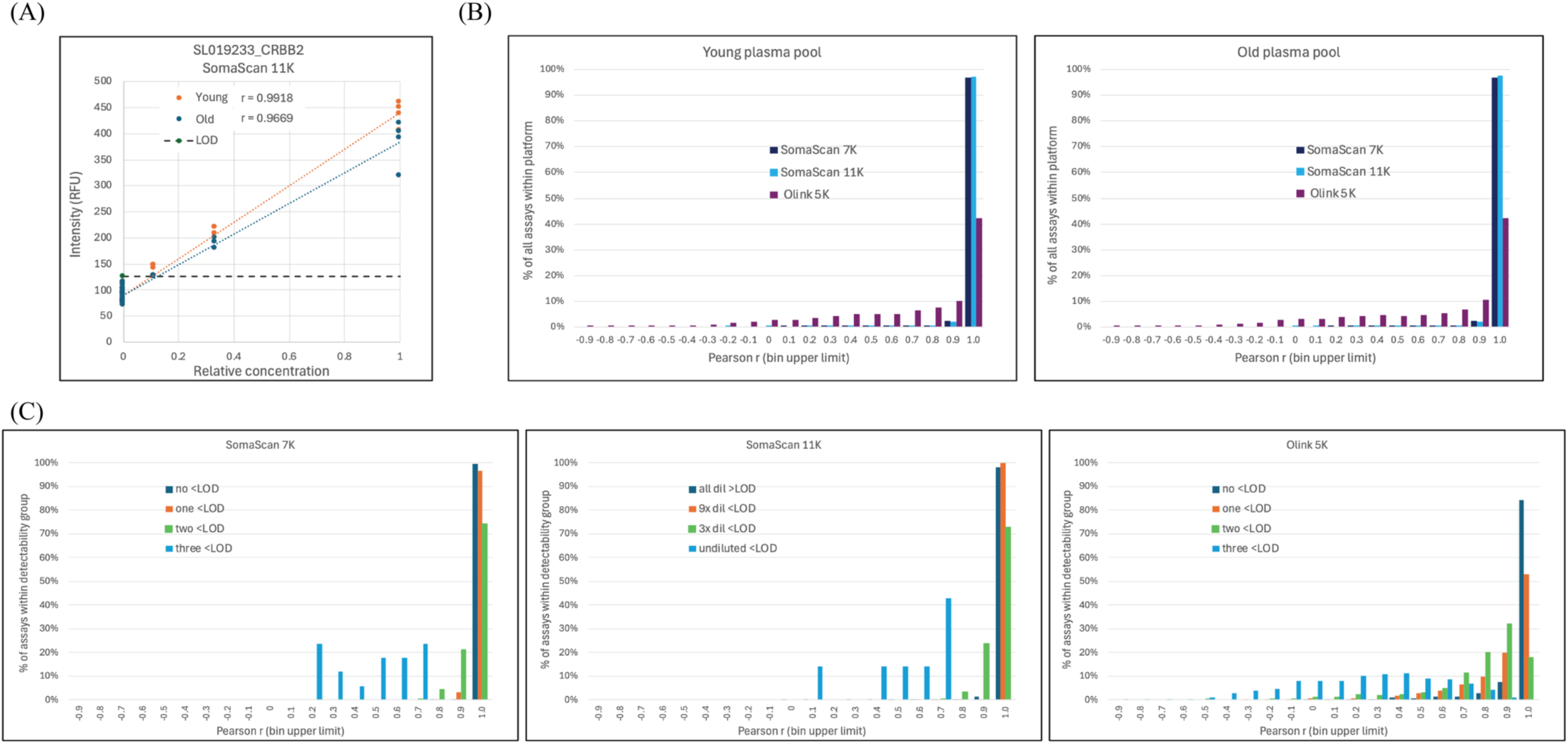
Dilution linearity in the SomaScan 7K, 11K, and Olink Explore 5K platforms for healthy plasma protein levels. (A) Example dilution linearity plot (SomaScan 11K data for two pooled healthy plasma samples and their 3x and 9x dilutions. eLOD = estimated limit of detection) (B) Distribution of Pearson correlation coefficient (r) of the plasma dilution data in the 3 platforms. (Total number of assays: SomaScan 7K: 7,288, SomaScan 11K: 10,776, Olink 5K: 5,416) (C) Linearity as function of detectability in the 3 platforms. Distribution of Pearson correlation coefficient (r) of the plasma dilutions for the 4 different detectability groups of the proteins. Protein assays were grouped depending on how many of the 3 plasma dilutions yielded signal below the eLOD.

**Figure S3:**
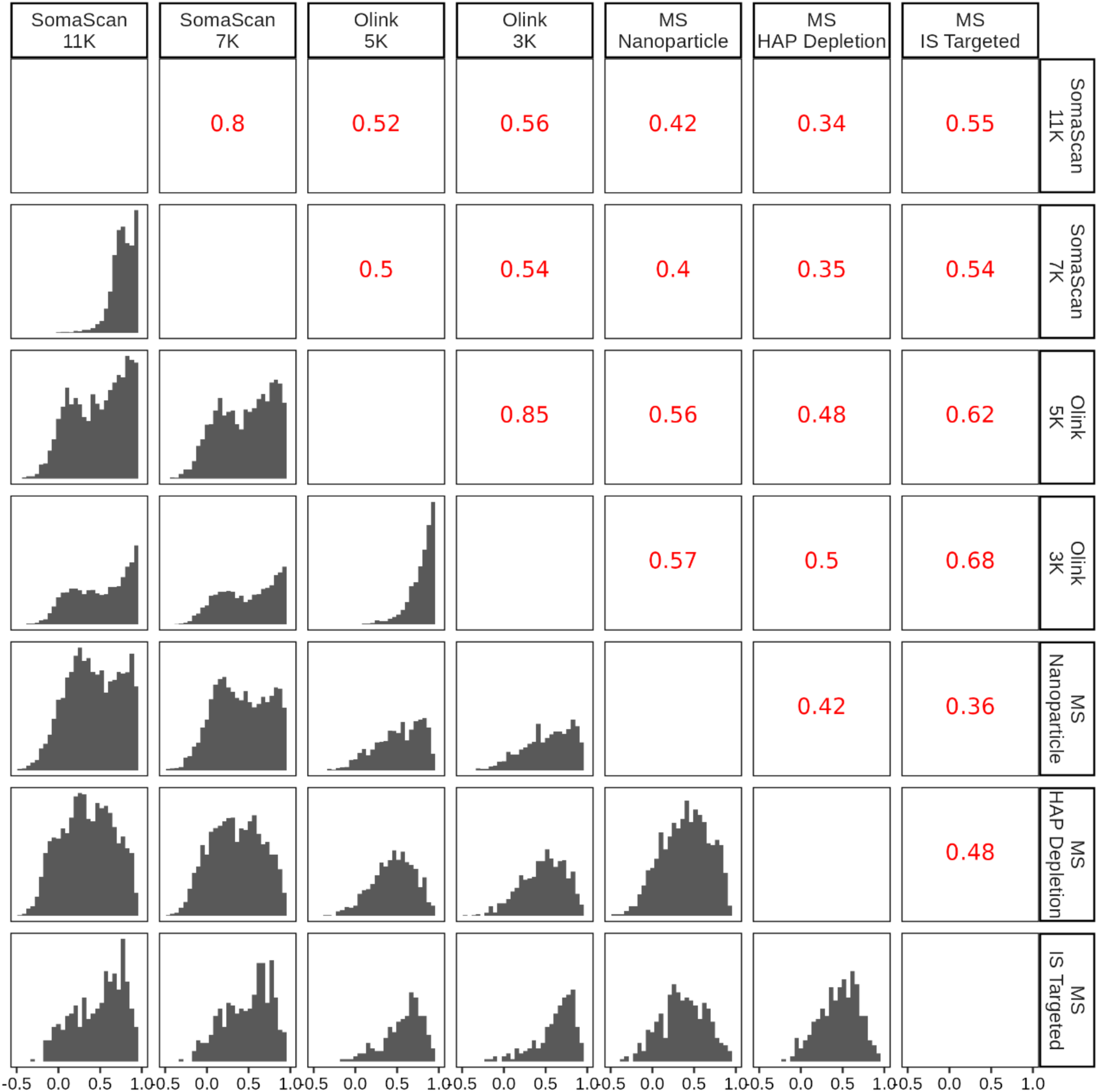
Distribution of Spearman Correlation Coefficients. Histogram showing the distribution of Spearman Rho correlation coefficient calculated for each protein. The median Spearman Rho correlation coefficient is shown in red. Data is restricted to subset of proteins that exceed eLOD and two-thirds data completeness.

**Figure S4:**
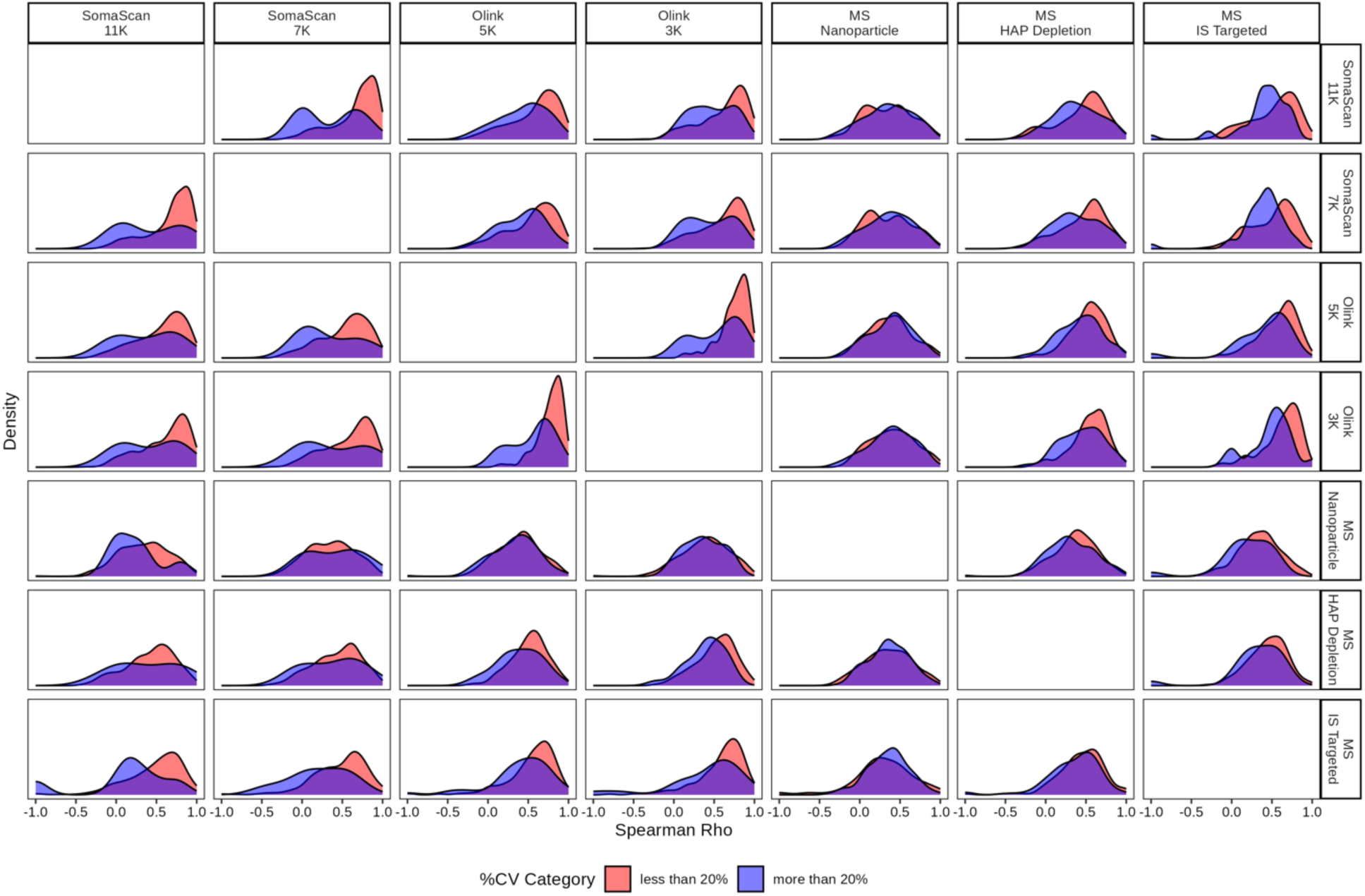
Density plots of the pairwise Spearman Rho correlation coefficients between platforms for the shared 254 proteins across all platforms. Each density plot is colored by binned technical cv of faceted column platform.

**Figure S5:**
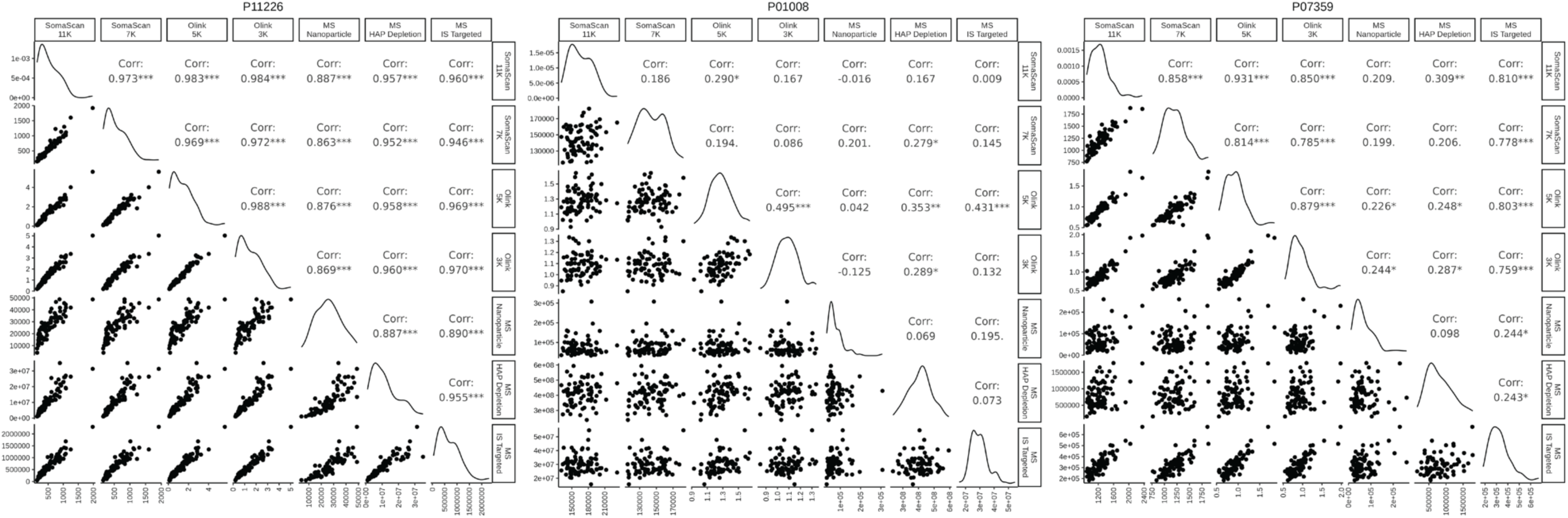
Inter-Platform Correlation of Selected Proteins. (A) P11226: Spearman correlation of protein P11226 across all platforms. (B) P01008: Spearman correlation of protein P01008 across all platforms. (C) P07359: Spearman correlation of protein P07359 across all platforms.

**Figure S6:**
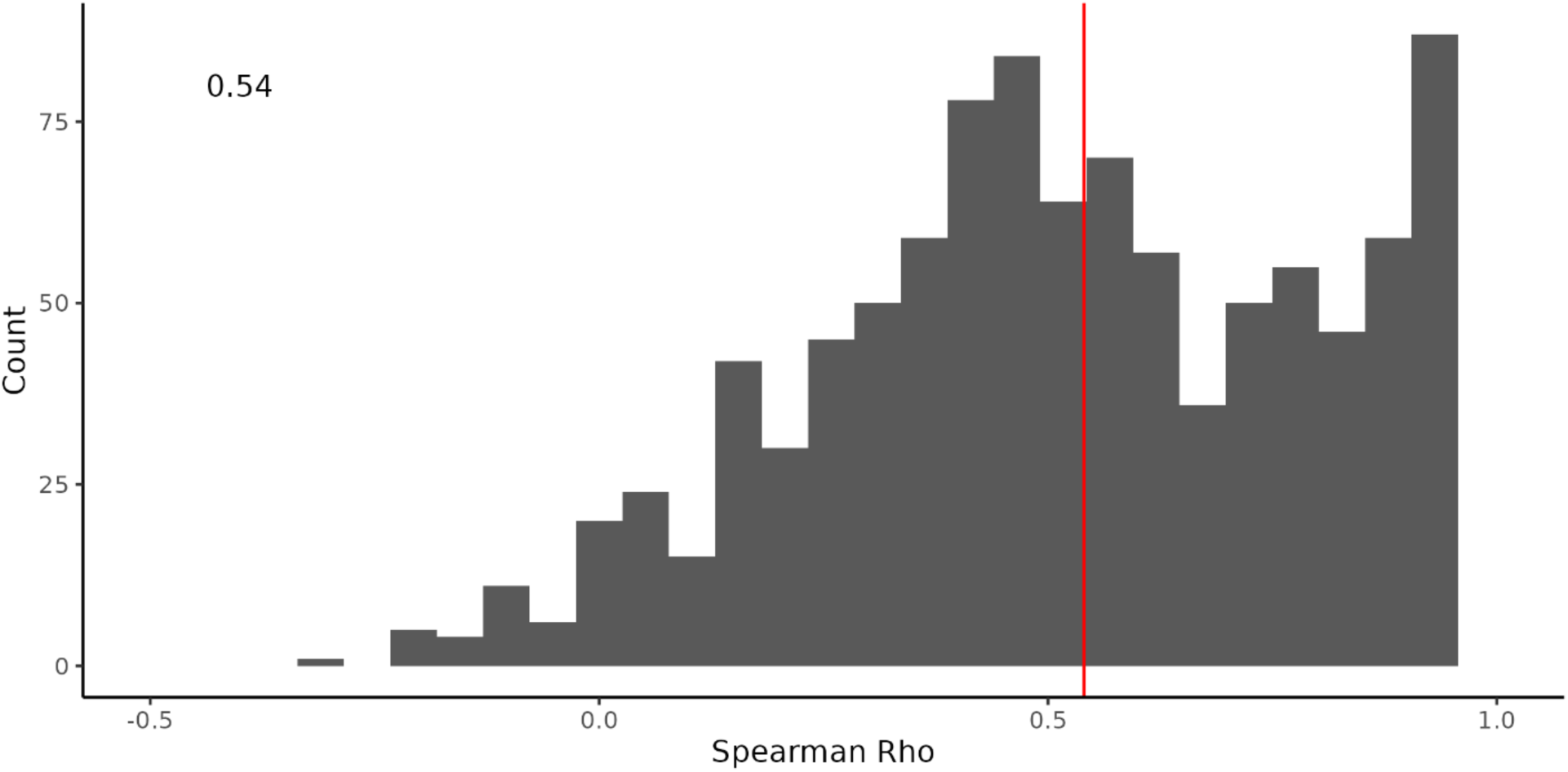
Correlation Between Aptamers Targeting the Same SOMAmer Target. Distribution of Spearman Rho correlations calculated between pairs of aptamers that target the same SOMAmer Target ID within the SomaScan 11K assay.

**Figure S7:**
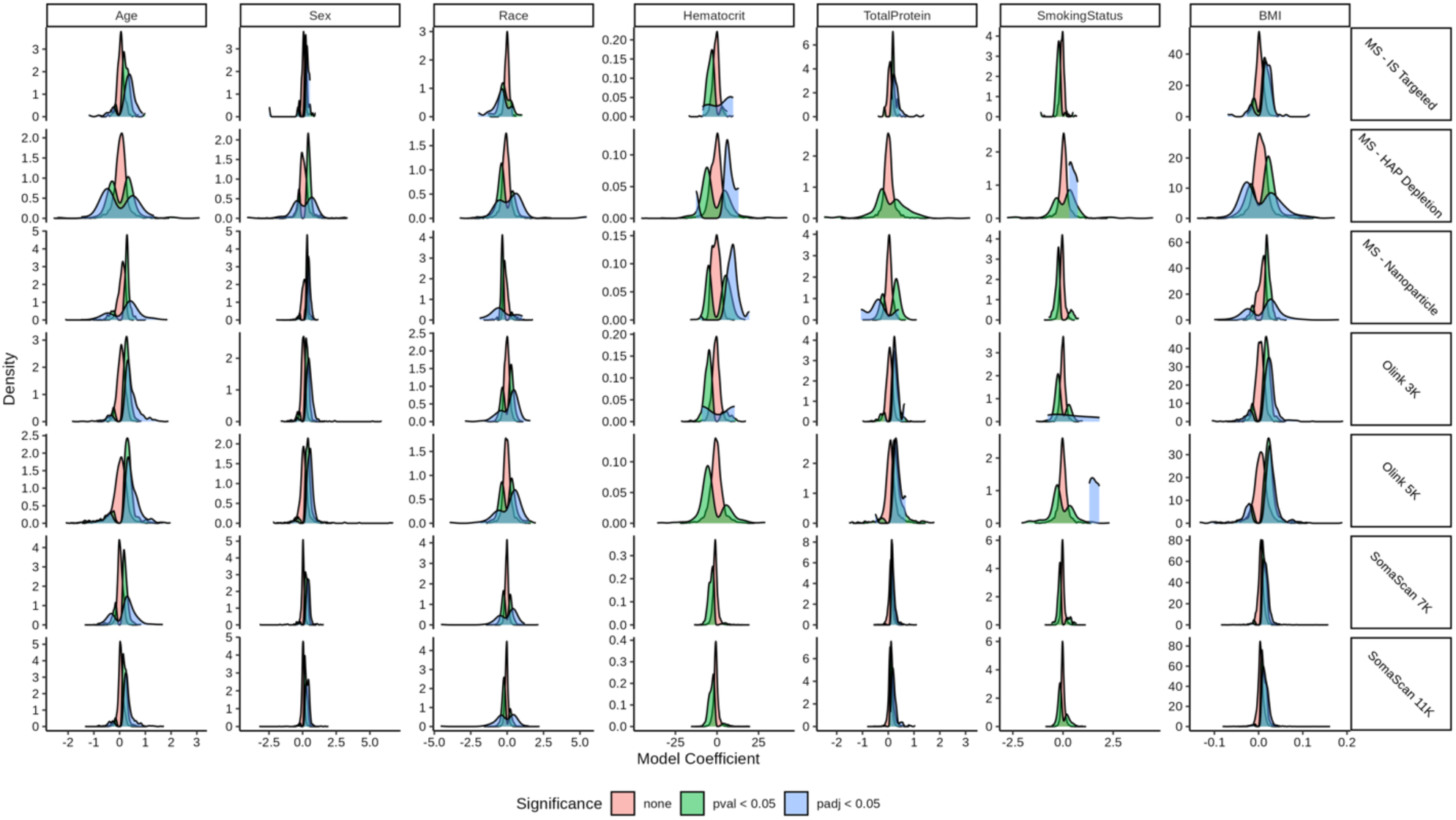
Distribution of Linear Model Coefficients by Significance. Each coefficient is colored according to its significance level. The distributions are shown separately for each factor and platform included in the model.

**Figure S8:**
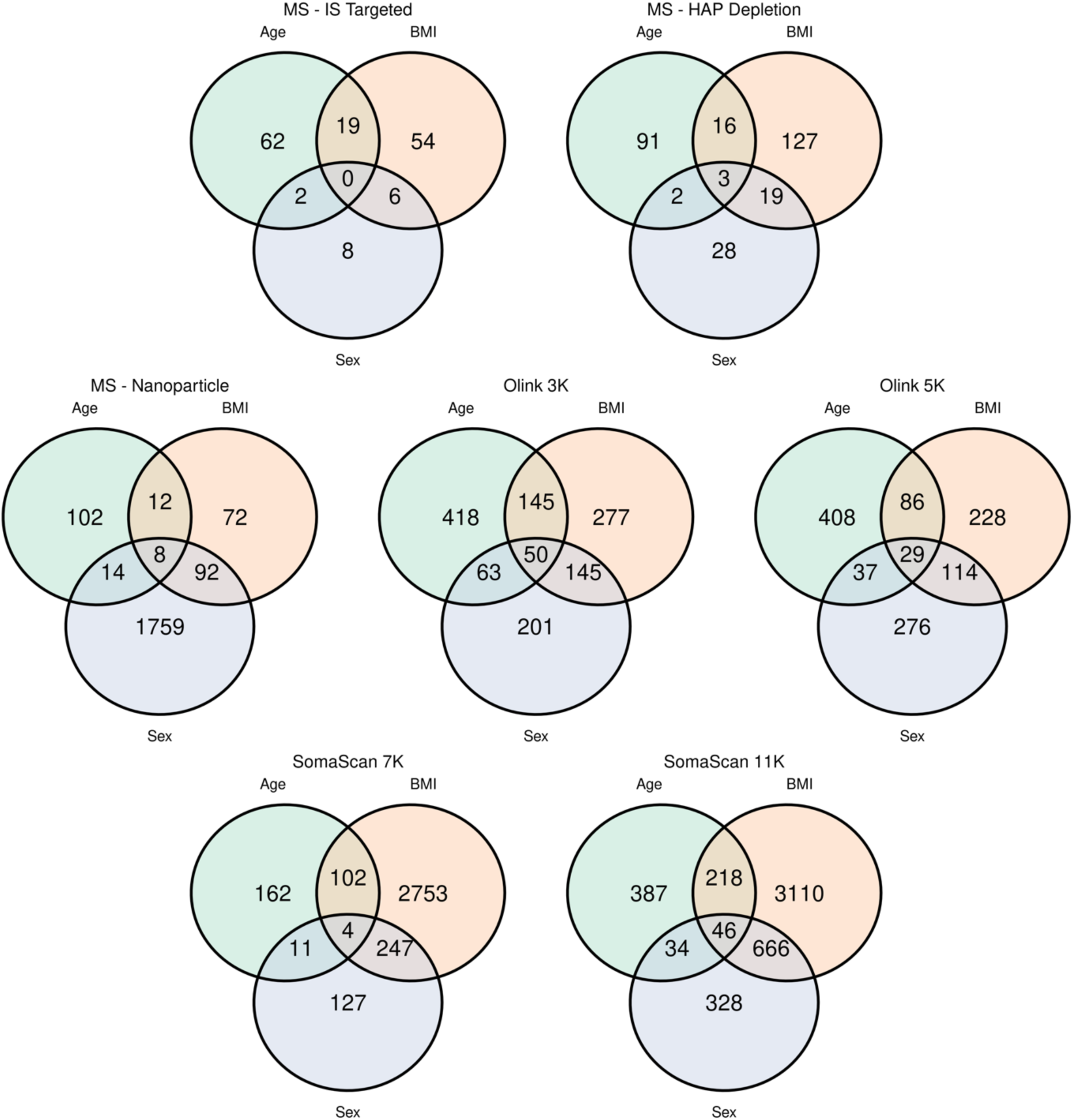
Overlap of Significant Biomarkers Across Platforms. Venn diagrams corresponding to significant (p-adj < 0.05) Age, BMI, and Sex markers across all platforms.

**Figure S9:**
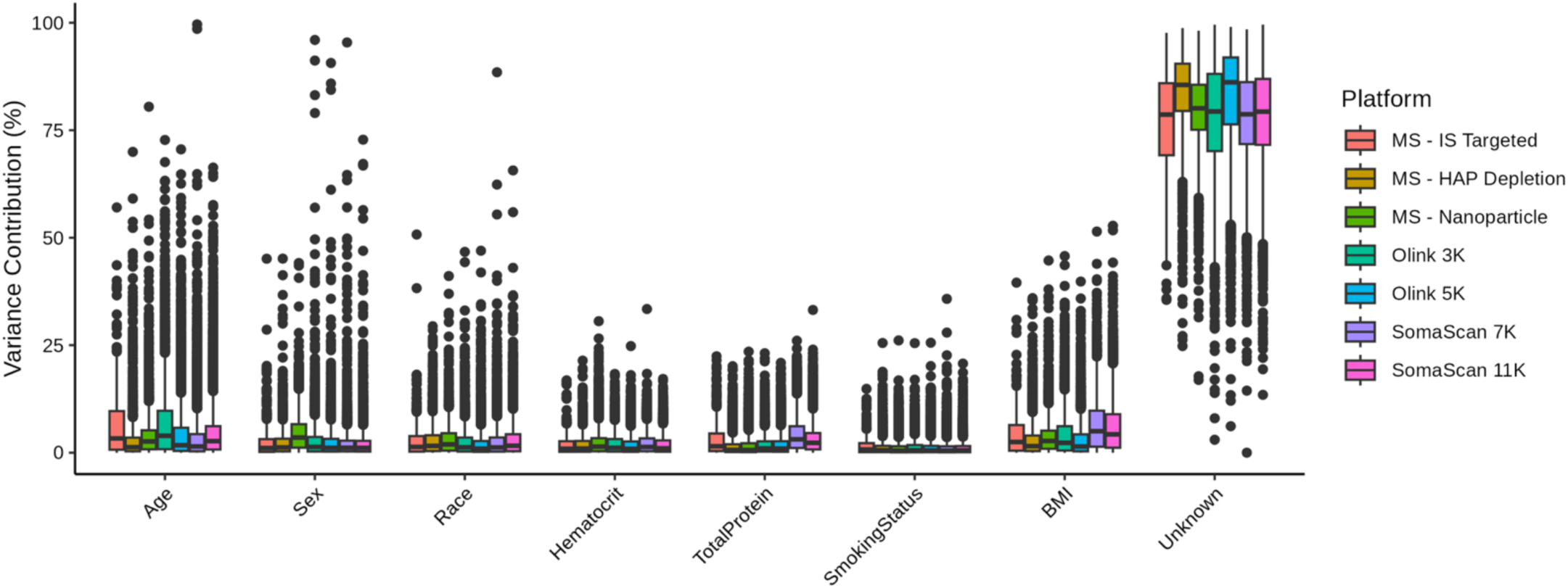
Variance Sources of Each Analyte. Percent of variance make-up identified from model parameters for each analyte

**Figure S10:**
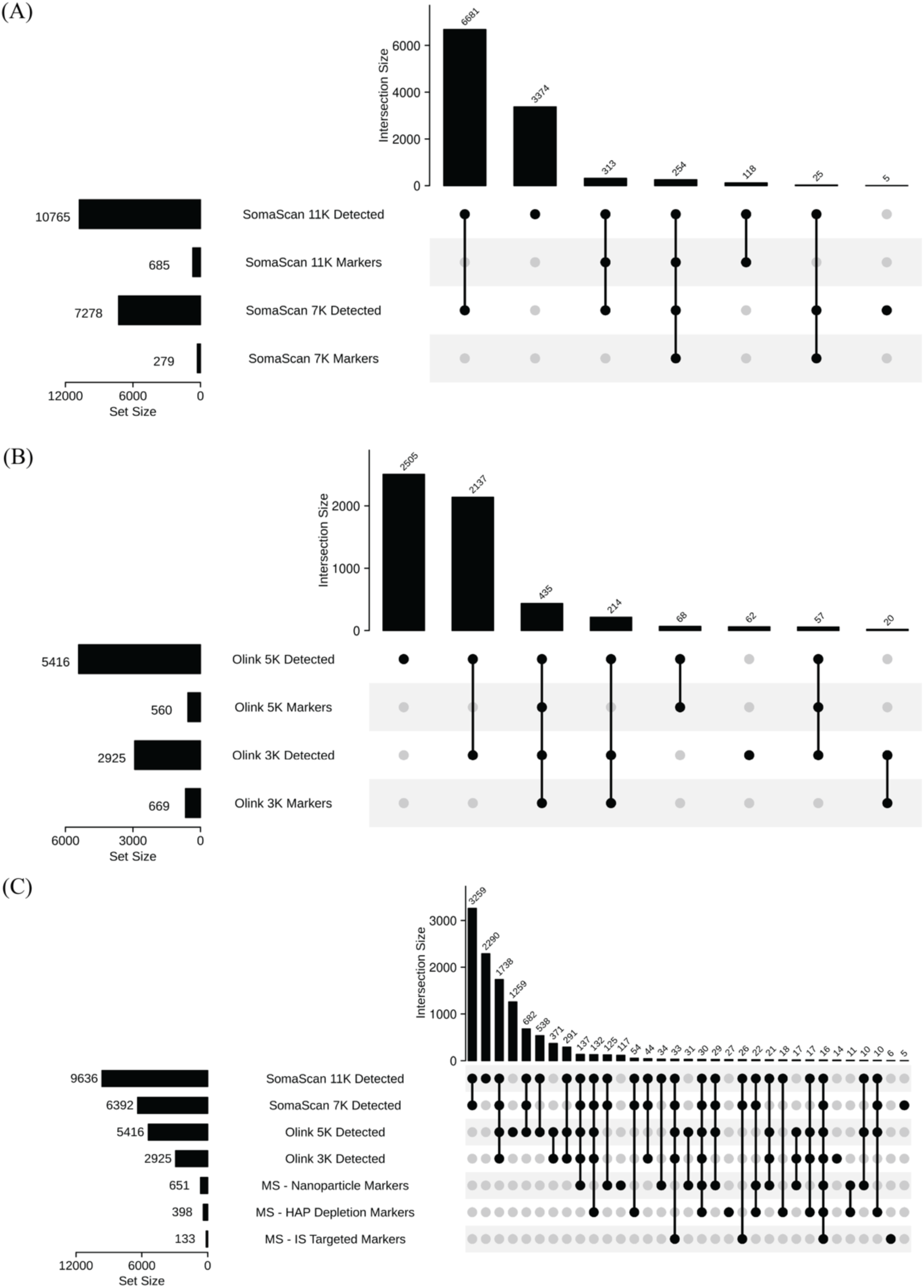
Age-Related Protein Markers Across Platforms. (A) UpSet plot of aptamers identified in SomaScan platforms and additionally identified as age markers (p-value < 0.05). (B) UpSet plot of proteins identified in Olink platforms and additionally identified as age markers (p-value < 0.05). (C) UpSet plot of proteins identified in SomaScan and Olink platforms and proteins identified age as markers in any Mass Spectrometry based platforms (p-value < 0.05).

## References

1. Yurkovich, J. T. & Hood, L. Blood is a window into health and disease. Clinical Chemistry vol. 65 1204–1206 Preprint at 10.1373/clinchem.2018.299065 (2019).

2. Anderson, N. L. & Anderson, N. G. The human plasma proteome: history, character, and diagnostic prospects. Molecular & cellular proteomics : MCP vol. 1 845–867 Preprint at 10.1074/mcp.R200007-MCP200 (2002).

3. Geyer, P. E., Holdt, L. M., Teupser, D. & Mann, M. Revisiting biomarker discovery by plasma proteomics. Mol Syst Biol 13, (2017).

4. Birhanu, A. G. Mass spectrometry-based proteomics as an emerging tool in clinical laboratories. *Clinical Proteomics* vol. 20 Preprint at 10.1186/s12014-023-09424-x (2023).

5. Deutsch, E. W. et al. Advances and Utility of the Human Plasma Proteome. Journal of Proteome Research vol. 20 5241–5263 Preprint at 10.1021/acs.jproteome.1c00657 (2021).

6. Ignjatovic, V. et al. Mass Spectrometry-Based Plasma Proteomics: Considerations from Sample Collection to Achieving Translational Data. Journal of Proteome Research vol. 18 4085–4097 Preprint at 10.1021/acs.jproteome.9b00503 (2019).

7. Geyer, P. E. et al. The Circulating Proteome─Technological Developments, Current Challenges, and Future Trends. J Proteome Res (2024) doi:10.1021/acs.jproteome.4c00586.

8. Lehallier, B. et al. Undulating changes in human plasma proteome profiles across the lifespan. Nat Med 25, 1843–1850 (2019).

9. Carrasco-Zanini, J. et al. Mapping biological influences on the human plasma proteome beyond the genome. Nat Metab (2024) doi:10.1038/s42255-024-01133-5.

10. Zaghlool, S. B. et al. Revealing the role of the human blood plasma proteome in obesity using genetic drivers. Nat Commun 12, (2021).

11. Geyer, P. E. et al. Plasma Proteome Profiling to Assess Human Health and Disease. Cell Syst 2, 185–195 (2016).

12. Zander, J. et al. Effect of biobanking conditions on short-term stability of biomarkers in human serum and plasma. Clin Chem Lab Med 52, 629–639 (2014).

13. Daniels, J. R. et al. Stability of the Human Plasma Proteome to Pre-analytical Variability as Assessed by an Aptamer-Based Approach. J Proteome Res 18, 3661–3670 (2019).

14. Cao, Z. et al. An Integrated Analysis of Metabolites, Peptides, and Inflammation Biomarkers for Assessment of Preanalytical Variability of Human Plasma. J Proteome Res 18, 2411–2421 (2019).

15. Lan, J. et al. Systematic Evaluation of the Use of Human Plasma and Serum for Mass-Spectrometry-Based Shotgun Proteomics. J Proteome Res 17, 1426–1435 (2018).

16. Tognetti, M. et al. Biomarker Candidates for Tumors Identified from Deep-Profiled Plasma Stem Predominantly from the Low Abundant Area. J Proteome Res 21, 1718–1735 (2022).

17. Keshishian, H. et al. Quantitative, multiplexed workflow for deep analysis of human blood plasma and biomarker discovery by mass spectrometry. Nat Protoc 12, 1683–1701 (2017).

18. Viode, A., et al. A Simple, Time-and Cost-Effective, High-Throughput Depletion Strategy for Deep Plasma Proteomics On Behalf of the IMPACC Network. https://tissues.jensenlab.org/ (2023).

19. Soni, R. K. Frontiers in plasma proteome profiling platforms: innovations and applications. Clin Proteomics 21, (2024).

20. Blume, J. E. et al. Rapid, deep and precise profiling of the plasma proteome with multi-nanoparticle protein corona. Nat Commun 11, (2020).

21. Sun, B. B. et al. Genomic atlas of the human plasma proteome. Nature 558, 73–79 (2018).

22. Bader, J. M., Albrecht, V. & Mann, M. MS-Based Proteomics of Body Fluids: The End of the Beginning. Molecular and Cellular Proteomics 22, (2023).

23. Van Eyk, J. E. & Sobhani, K. Precision Medicine: Establishing Proteomic Assessment Criteria from Discovery to Clinical Diagnostics. Circulation vol. 138 2172–2174 Preprint at 10.1161/CIRCULATIONAHA.118.036781 (2018).

24. Suhre, K. et al. Nanoparticle enrichment mass-spectrometry proteomics identifies protein-altering variants for precise pQTL mapping. Nat Commun 15, (2024).

25. Katz, D. H., et al. *Proteomic Profiling Platforms Head to Head: Leveraging Genetics and Clinical Traits to Compare Aptamer-and Antibody-Based Methods*. *Sci. Adv* vol. 8 https://www.science.org (2022).

26. Raffield, L. M. et al. Comparison of Proteomic Assessment Methods in Multiple Cohort Studies. Proteomics 20, (2020).

27. Candia, J. et al. Assessment of Variability in the SOMAscan Assay. Sci Rep 7, (2017).

28. Rooney, M. R. et al. Plasma proteomic comparisons change as coverage expands for SomaLogic and Olink. Preprint at 10.1101/2024.07.11.24310161 (2024).

29. Hendricks, N. G. et al. An inflection point in high-throughput proteomics with Orbitrap Astral: analysis of biofluids, cells, and tissues. Preprint at 10.1101/2024.04.26.591396 (2024).

30. Bhowmick, P., Roome, S., Borchers, C. H., Goodlett, D. R. & Mohammed, Y. An Update on MRMAssayDB: A Comprehensive Resource for Targeted Proteomics Assays in the Community. J Proteome Res 20, 2105–2115 (2021).

31. Eldjarn, G. H. et al. Large-scale plasma proteomics comparisons through genetics and disease associations. Nature 622, 348–358 (2023).

32. Clemente-Suárez, V. J. et al. The Role of Adipokines in Health and Disease. Biomedicines vol. 11 Preprint at 10.3390/biomedicines11051290 (2023).

33. Hellström, L., Wahrenberg, H., Hruska, K., Reynisdottir, S. & Arner, P. Mechanisms behind gender differences in circulating leptin levels. J Intern Med 247, 457–462 (2000).

34. Shah, N. R. & Braverman, E. R. Measuring adiposity in patients: The utility of body mass index (BMI), percent body fat, and leptin. PLoS One 7, (2012).

35. Reed, E. R. et al. Cross-platform proteomics signatures of extreme old age. Preprint at 10.1101/2024.04.10.588876 (2024).

36. Rauzier, C., Lamarche, B., Tremblay, A. J., Couture, P. & Picard, F. Associations between insulin-like growth factor binding protein-2 and lipoprotein kinetics in men. J Lipid Res 63, (2022).

37. Llobet, M. O., Johansson, Å., Gyllensten, U., Allen, M. & Enroth, S. Forensic prediction of sex, age, height, body mass index, hip-to-waist ratio, smoking status and lipid lowering drugs using epigenetic markers and plasma proteins. Forensic Sci Int Genet 65, (2023).

38. Wang, H. et al. Quantitative iTRAQ-based proteomic analysis of differentially expressed proteins in aging in human and monkey. BMC Genomics 20, (2019).

39. Sathyan, S. et al. Plasma proteomic profile of age, health span, and all-cause mortality in older adults. Aging Cell 19, (2020).

40. Sun, B. B. et al. Plasma proteomic associations with genetics and health in the UK Biobank. Nature 622, 329–338 (2023).

41. Ferdosi, S. et al. Engineered nanoparticles enable deep proteomics studies at scale by leveraging tunable nano-bio interactions. (2022) doi:10.1073/pnas.

42. Huang, T., et al. Protein Coronas on Functionalized Nanoparticles Enable Quantitative and Precise Large-Scale Deep Plasma Proteomics. *bioRxiv* (2023) doi:10.1101/2023.08.28.555225.

43. Wang, B. et al. Comparative studies of genetic and phenotypic associations for 2,168 plasma proteins measured by two affinity-based platforms in 4,000 Chinese adults. Preprint at 10.1101/2023.12.01.23299236 (2023).

44. Rooney, M. R. et al. Comparison of Proteomic Measurements Across Platforms in the Atherosclerosis Risk in Communities (ARIC) Study. Clin Chem 69, 68–79 (2023).

45. Candia, J. et al. Variability of 7K and 11K SomaScan Plasma Proteomics Assays. J Proteome Res 23, 5531–5539 (2024).

46. Dammer, E. B. et al. Multi-platform proteomic analysis of Alzheimer’s disease cerebrospinal fluid and plasma reveals network biomarkers associated with proteostasis and the matrisome. Alzheimers Res Ther 14, (2022).

47. Emilsson, V. et al. Co-regulatory networks of human serum proteins link genetics to disease. Science *(1979)* 361, 769–773 (2018).

48. Argentieri, M. A. et al. Proteomic aging clock predicts mortality and risk of common age-related diseases in diverse populations. Nat Med (2024) doi:10.1038/s41591-024-03164-7.

49. Coenen, L., Lehallier, B., de Vries, H. E. & Middeldorp, J. Markers of aging: Unsupervised integrated analyses of the human plasma proteome. Frontiers in Aging 4, (2023).

50. Pham, T. V., Henneman, A. A. & Jimenez, C. R. Iq: An R package to estimate relative protein abundances from ion quantification in DIA-MS-based proteomics. Bioinformatics 36, 2611–2613 (2020).

51. Nevola K, S. M. G. J. F. S. C. C. P. P. Z. B. S. M. D. K. K. A. C. L. C. K. OlinkAnalyze: Facilitate analysis of proteomic data from Olink. https://github.com/Olink-Proteomics/OlinkRPackage https://www.olink.com/patents/. (2024).

52. R Core Team. R: A language and environment for statistical computing (R Foundation for Statistical Computing). Preprint at https://www.R-project.org/ (2023).

53. Alexa A & Rahnenfuhrer J. topGO: Enrichment Analysis for Gene Ontology. (2024).

54. Yu, G. & He, Q.-Y. ReactomePA: an R/Bioconductor package for reactome pathway analysis and visualization. Mol Biosyst 12, 477–479 (2016).

55. Wu, T. et al. clusterProfiler 4.0: A universal enrichment tool for interpreting omics data. Innovation 2, (2021).

56. Yu, G., Wang, L. G., Han, Y. & He, Q. Y. ClusterProfiler: An R package for comparing biological themes among gene clusters. OMICS 16, 284–287 (2012).

57. Wickham, H. Ggplot2: Elegant Graphics for Data Analysis. (Springer-Verlag New York, 2016). doi:10.1007/978-3-319-24277-4.

58. Gu, Z., Eils, R. & Schlesner, M. Complex heatmaps reveal patterns and correlations in multidimensional genomic data. Bioinformatics 32, 2847–2849 (2016).

59. Gu, Z. Complex heatmap visualization. iMeta 1, (2022).

